# A workcell 1.0 for programmable and controlled operation of multiple fluidic chips in parallel

**DOI:** 10.1101/2023.04.16.536594

**Authors:** Chuanfang Ning, Gabriel Bunke, Simon Lietar, Lukas van den Heuvel, Amir Shahein

## Abstract

We developed a versatile lab-on-chip (LOC) workcell that enables the design and automatic execution of experiments on LOC devices, improving how we establish, optimize, and productionalize LOC processes. Key features include direct docking and cooling of native laboratory tubes, programmable reagent mixing and dilutions, parallel operation of multiple chips, precise flowrate and pressure control, clogging detection and response, programmable microscope control, chip temperature regulation, and scheduled cleaning. All functionality is controlled seamlessly from an easy-to-write protocol file, and based on extensible hardware and software infrastructures to promote community development. To showcase the platform’s use and versatility, we demonstrate a series of 5 different automated experiments at varying levels of complexity, executed across both Quake-valve and droplet microfluidic systems. In particular, the workcell was instructed to map the parameter regime that generates viable droplets, to allow a user to select diameters and production frequencies of interest for single bacterial cell encapsulation. Furthermore, three out of three days in a row, the platform successfully performed a complex 15.5h long experiment, integrating in a single automated protocol the full core workflow required by a typical protein-characterization lab: protein expression, purification, dilution generation, and quantitative binding characterization (generating 55296 images in the process). Experiments conducted through the workcell are easier to set up, offer increased control over experiment conditions and parameters, and can be heavily parallelized.

## Introduction

Compared to other research tools for liquid handling and experimentation, lab-on-chip (LOC) devices carry numerous advantages across the many different contexts in which they are used. Typical mentions include higher achievable throughputs, reduced reagent consumption, or the enabling of experiments that are not otherwise possible, like trapping and thoroughly characterizing large numbers of single cells ^1–4^, next-generation sequencing^5, 6^, or organ-on-chips ^7^. On the other hand, interfacing with LOC systems is often done manually, and significant effort can be necessary to robustly integrate different LOC modules ^8^, to establish reproducibility, and to translate processes to production and scale.

Despite the past 30 years of research^9, 10^, the prevailing belief is that microfluidics is in its adolescence and has yet to materialize its potential ^9, 11–16^. While one can quickly find a well-published, intheory better microfluidic counterpart to most common laboratory procedures ^2, 12, 17–23^, and microfabrication methods usually allow for the possibility to integrate these compact modules inline ^8^, we still have not realized the longstanding goal of routine integration of modules to compose complex, multi-step workflows. Notable LOC-like counterexamples include integrated protein production, purification, and characterization workflows ^24, 25^, single-cell isolation, stimulation, and characterization ^1–4^, NGS prep ^26–28^, and next-generation sequencing ^5, 6^. Perhaps more striking yet, is the small proportion of solutions validated in fluidics research that have resulted in mainstream adoption ^9, 12^, and the limited number of cases where companies have successfully productionalized fluidic chips for industrial applications. Interesting counterexamples, however, exist at the technological core of large companies, including Abcellera, Illumina, 10X genomics, Oxford Nanopore, Twist Bioscience, Carterra, Fluidigm, and Berkeley lights, with growing numbers of programs inside of big pharma ^29^.

While such delays in system integration and adoption are likely a result of multiple causes, we share the belief that one important reason is insufficient supporting infrastructure and technologies to facilitate lab-on-chip process development^12, 15^. In particular, how experiments are designed and executed on LOC devices typically does not adhere to engineering practices ensuring abstraction in experiment design, software and hardware modularity, good error-handling, standardization, and parallelization of experimentation. Laboratory personnel rely either on manual implementation, custom hardware and software that carries limited features and lacks versatility, or incomplete proprietary solutions offering limited design of and control over complex experiments.

Furthermore, there is currently little open-source tooling available that is useful for taking chip-based processes into production^12, 30^. Whereas in the research lab, the objective is often to establish a process that is novel and delivers a valuable outcome, in production, the objective is to ensure that this outcome is achieved more reproducibly and at scale. This can be challenging when working with sensitive biology and as the complexity and novelty of a laboratory process increases. To establish reproducibility, put simply it is necessary to mitigate sources of variability, which may involve standardizing process protocols (experiment steps) and the generation of process inputs (reagent formulations and dilutions, chip fabrication and loading), as well as developing control over process parameters (flowrates, pressures) and environmental conditions (humidity, reagent and reaction temperatures). Scale can be achieved both by redesigning at a chip-level, or through automation and parallelization of chip operation.

In order to improve how the scientists in our group work with lab-on-chip processes, we developed an LOC workcell that through an accessible language enables its users to conveniently design experiments, which are then run automatically at a high-level of standardization and control, while allowing for realtime experiment management. A user selects or designs the experiment of interest using the Automancer software ^31^, screws the necessary aliquot tubes into the workcell’s reagent hub, and clicks run to begin. By integrating plug-and-play with standard laboratory freezer tubes, and by conducting programmable reagent mixing and dilutions as specified in the protocol file, we remove entirely the need for manual pipetting or reagent loading.

At the reagent hub, reagents are cooled to a specified temperature by a custom cooling system, and shuttled from their tubes to be introduced into the chips at the appropriate time in the experiment. Reagents travel as fluid packets shepherded by air using a Load, Advance, Introduce, Reset (LAIR) flow protocol, and can be introduced into target chips at either a precise flowrate (by way of a custom flowrate controller) or a precise pressure. Air is prevented from entering a chip’s functional region. With little difference in experiment setup or protocol file, the workcell can implement the experiment on either a single chip, or on multiple chips in parallel to increase process throughput. When running multiple chips, they can be dynamically coupled or decoupled as necessary during experiments, to drive a given reagent to all of the chips at once or to enable different protocol steps to be run on different chips, respectively.

Chip imaging is automated through programmatic control of a Nikon Eclipse Ti2 directly from the user’s protocol file, with the ability to specify a cloud storage location (e.g. S3 bucket in Amazon Web Services (AWS)) as destination for captured images. The workcell uses a heating controller to set chip temperatures for incubation steps and to improve environment control. We also demonstrate how automated cleaning occurs through the flushing of any reused hardware and lines with cleaning solution at the end of every protocol, further improving standardization.

We placed emphasis on the tracking of process parameters, like chip flowrates, line pressures, clogging events, chip and reagent temperatures, which can be saved for part or all of an experiment, helping the lab to identify events that may compromise results and eventually helping to implement quality control. Experiment execution can also respond live to sensor feedback, enabling automatic event detection and response to prevent system failures, which we use to for instance detect and respond to chip-clogging. In the event of partial clogging, the control architecture will ensure flowrate robustness and continued operation. If the degree of clogging is too great, such that compensatory pressures for flowrate stability threaten chip delamination, then the experiment is instead automatically frozen into the nearest userlabeled safe state (e.g. valves closed and pressures off, until the user intervenes). All of the hardware and functionality is controlled from a single, versionable and shareable protocol file that is easy to write and read by researchers not having any prior programming experience.

To highlight the use and versatility of our platform, we demonstrate a series of 5 different automated experiments that take advantage of different system features, and execute across both Quake-valve and droplets chips. We demonstrate on-chip protein expression and purification, dilution generation by inlet valve pulse-width modulation paired with on-chip multiplexing for routing, and a simple binding assay. We also demonstrate a 15.5 hour-long quantitative binding characterization experiment generating 18432 images, where the proteins being characterized are expressed and purified directly on-chip prior to being characterized against 6 concentrations of their binding partner, and which ran successfully three out of three experimental days in a row. Using droplet microfluidics we demonstrate an automated parametersearch experiment for optimization. The workcell iteratively varies the flowrates of continuous and dispersed phases across 105 different conditions, and captures an image of the droplets at each condition to map the parameter regime that generates viable droplets and to allow the user to select accessible droplet diameters and production frequencies of interest. The workcell then encapsulates single bacterial cells at these specified diameters.

We open-source all hardware and software required to build this platform. Since we plan to grow the workcell in our own lab, we emphasized modular and scalable hardware and software infrastructures in development, which should allow other researchers to build a version that suits their function and throughput requirements. We expect that the full workcell or particular modules will be useful for other labs aiming to improve how they develop, optimize, and productionalize lab-on-chip processes.

## Workcell design

### Design objectives

The LOC workcell was designed to satisfy several objectives related to scale, reproducibility, and programmability. We designed the platform to be used in a research-lab environment, for improving and scaling the characterization of LOC devices and assay versions in development, for device optimization, and for high-throughput data generation as an LOC process becomes stable and enters production. We were particularly interested in satisfying the operation of more complex LOC systems relying on active fluid routing, where abstraction of experiment design and comprehensive control may provide the most value.

To lay the foundation for our objectives we pursued full platform automation. To increase throughput, we sought to operate multiple chips in parallel from the same input reagent tubes, while allowing for independent addressability of chips to increase the number of possible experiment variations that can be tested at once. In addition to a decrease in labor cost through automation, we sought to reduce reagent expenditure, and to minimize the initial experiment setup time. We further targeted the mitigation of sources of variability through tracking and control over process parameters and environmental conditions, by removing the need for manual and inconsistent reagent mixing, dilutions or loading, as well as through leveraging automation for protocol standardization and text-based protocols for versioning. Programmability means that hardware should be able to be reconfigured based on a change in instruction-set (protocols, configuration files), and that these instructions should be able to be designed conveniently by any user. Additionally, we desired to make other process functionality, like flowrates, fluid-routing, responses to critical events like clogging, and reagent dilutions as freely programmable as possible, to hand over more control and versatility to the user. The high-level design, as well as the specific starter pack of modules and functionality that we include should be versatile enough to satisfy a range of LOC applications, such that these different processes can be implemented through a similar, standardized platform and workflow. Furthermore, the hardware and software should be modular and easily extensible, to minimize cost for supporting applications not covered by the initial design (in particular, where different actuators are preferred for fluid routing, or detectors for characterization).

### Hydra fluid control system

The developments that comprise the workcell consist of a combination of modules on-chip and peripheral hardware, together with software and fluid handling protocol (see Figure 1, and the corresponding 3D CAD animation in Supplementary Video 1). We built the Hydra reagent control system to maintain reagents at a specified temperature throughout an experiment and to deliver reagents to the chips (Figure 1A). The design is described in more detail in Supplementary Section 1.1, with CAD files available in the associated Github repository. The user simply takes cryo aliquot tubes from the freezer and screws them directly into the Hydra’s pressure head, which mates with a custom cooling block (Figure 2E–F). No manual fluid manipulations are required, even concerning the mixing or generation of dilutions of reagents. A given reagent can be programmatically selected to travel out of the pressure head through one of two connecting rotary distributor valves at a specified pressure or flowrate. Distributor valves are electronic many-to-1 valves, together selectively routing up to 24 input tubings (of pressurized reagent) to 2 output tubings. Maintaining integrity of reagents was important, and so our design ensures that reagents do not come into contact with any internal surfaces of the pressure head during ejection, apart from a standard biocompatible tubing of choice such as PTFE or tygon. The two outputs from the two rotary valves can either feed directly into two inlets of a chip, or can each be split into *n* lines for parallelized chip-running of *n* chips (we have tested up to *n*=3, although we expect this can be increased further). When parallelizing chip-running, minimal changes are required relative to the single chip case. Simply the rotary valve output line must be split, more reagent should be contained in the source tubes, and different functions should be imported into the experiment protocol file. In order to transfer a reagent from its tube into the functional region of the chip, we drive a fluid packet with air using a flow protocol that we refer to as Load Advance Introduce Reset (Figure 2A–D).

**Figure 1:**
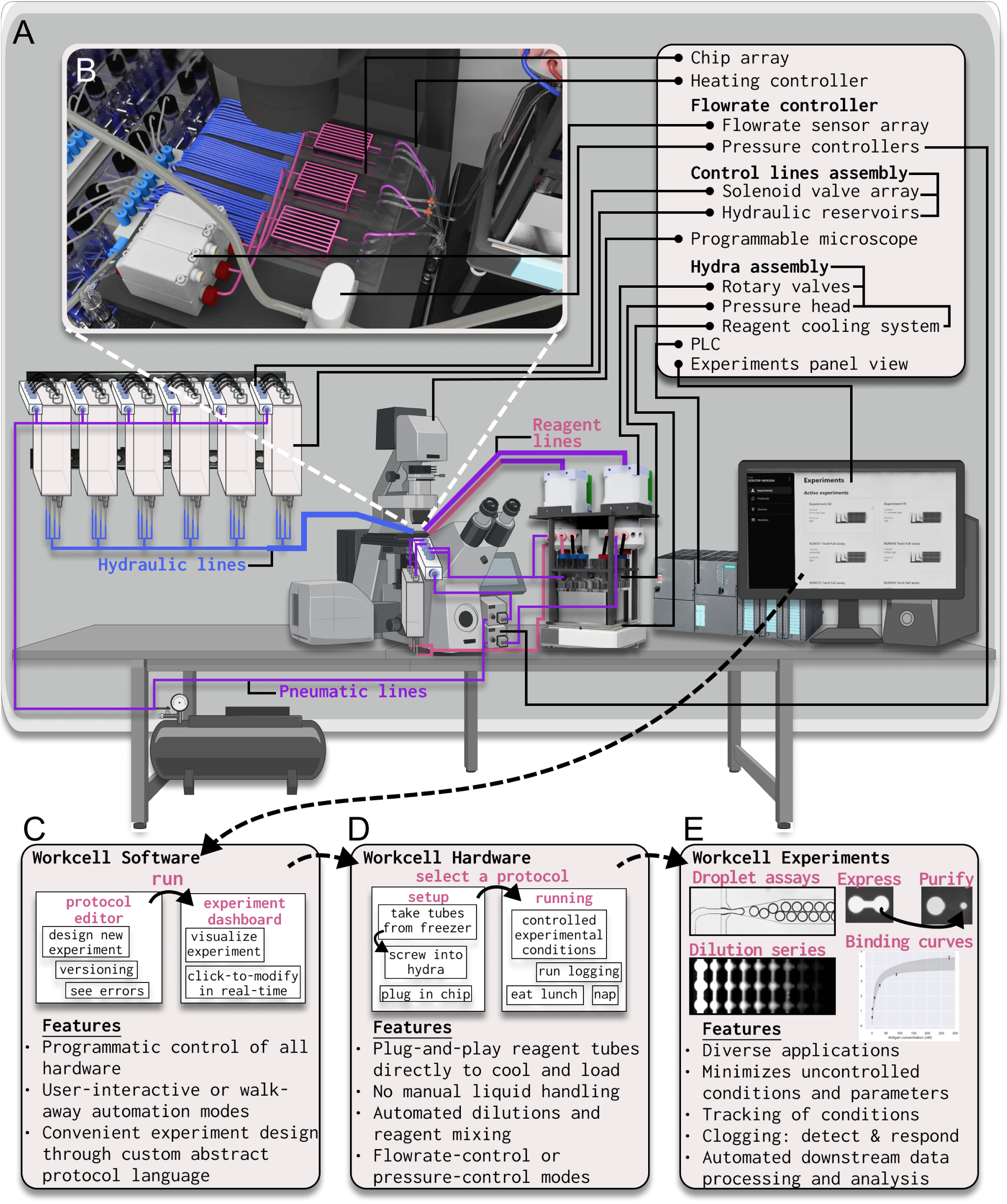
Overview of the workcell. Caption on the next page. Figure 1: **(A)** The workcell is composed of a set of on-chip modules, peripheral hardware, software, and fluid handling protocol. An overview of some aspects of the workcell is available as a 3D CAD animation in Supplementary Video 1. Peripheral hardware includes a hydra assembly (fluid control system), where standard laboratory freezer tubes are docked, cooled, and pressurized to travel by Load Advance Introduce Reset cycles (LAIR) through reagent lines to an array of chips **(B)**. The workcell also includes an automated Quake-valve control system, flow and pressure controllers, a programmable microscope, a heating controller, and a programmable logic controller to integrate and control peripheral hardware. **(C)** High-level control software (automancer ^31^) runs on a desktop application with a user-interface, while latency-critical programs are implemented directly on a programmable logic controller. Experiments are designed using a custom abstract protocol language. They can be versioned (see Supplementary Figure 6), and managed in real-time. **(D)** Workcell hardware enables simple experiment setup, high levels of standardization and precise control. **(E)** The workcell can execute diverse experiments.

### LAIR flow protocol

During the Load stage, a rotary valve switches to the position of the desired reagent, and the reagent is then pressurized to travel through the tubing connecting to the rotary valve, and into the output line travelling from the rotary valve to the chip (Figure 2A). The amount of liquid that is loaded into the output line will be selected based on the flow cycle’s future Introduce stage, with a small surplus that will be cleared in the Reset stage to account for error (see Table 1 for volume and timing characterizations, or Supplementary Figure 1). On our workcell, we typically use Load stages of between 0.5s and 10s depending on the loaded volume.

**Figure 2:**
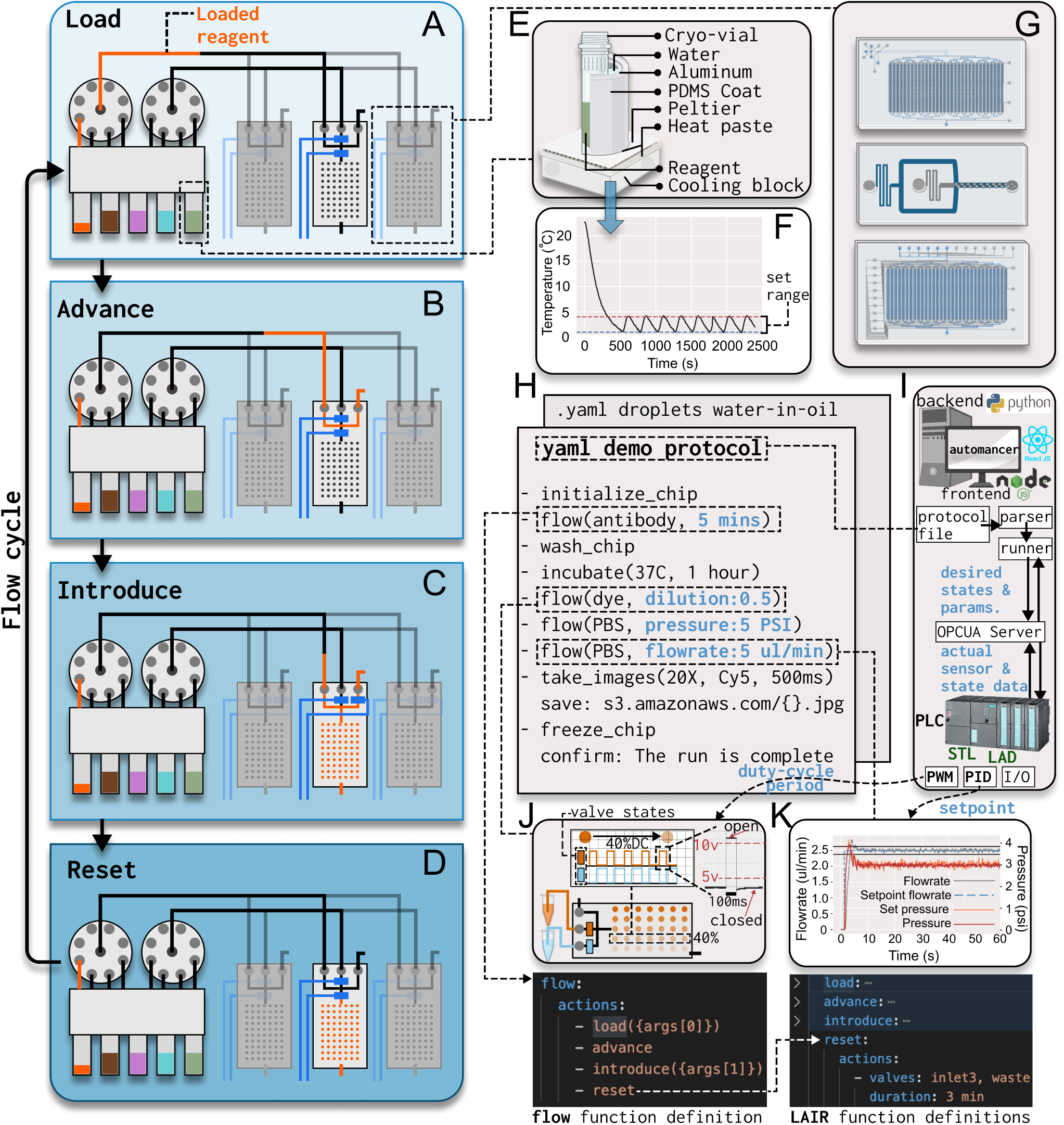
LAIR flow protocol, software architecture, demo protocol, and system characterizations. Caption on the next page. Figure 2: **(A-D)** Schematic depicting the Load, Advance, Introduce, and Reset stages of the flow protocol that is used to move reagents from their tubes to an array of chips an arbitrary distance away, with low dead-volume. **(E)** Layout showing the different layers of the cooling block, which the tubes sit in while they are docked to the pressure-head. **(F)** Characterization of the cooling system. Temperature readings are generated by a high-precision resistance temperature detector (RTD, Pt-100) which sits in water inside of a dummy laboratory tube. **(G)** Chips that were used on the workcell in this study. Chip from Figure 5A, C, E (top), chip from Figure 5D (middle), chip from Figure 5B (bottom). **(H)** Demo protocol for controlling the workcell to operate chips (functional syntax is similar in simplicity). Hierarchical function definitions are shown below for “flow”, which calls “Load”, “Advance”, “Introduce”, and “Reset” functions. Functions are imported at the beginning of a protocol file. **(I)** Schematic overview of software architecture. The automancer software ^31^ communicates by OPCUA to software and hardware running on the PLC. **(J-K)** Depictions of programs running on the PLC. **(J)** Depiction of inlet-valve pulse-width modulation (PWM) used to automate reagent mixing and on-chip dilutions. Automancer communicates to turn on PWM-mode for a specific channel, and supplies a period, and duty-cycle to the PLC, to select a specific dilution level or mixing ratio. Schematic waveforms are shown for a 40% duty cycle used to dilute orange reagent (top). An example pulse from the PLC is shown as characterized using a logic analyzer (top right). More extensive PWM characterization is shown in Supplementary Figure 3. **(K)** Characterization of the flowrate controller. Automancer directs the PLC to operate in flowrate-control mode, and supplies a flowrate setpoint. A characterization across a wide range of flowrates is shown in Supplementary Figure 2, and stability across resistance perturbations in Figure 4.

**Table 1:**
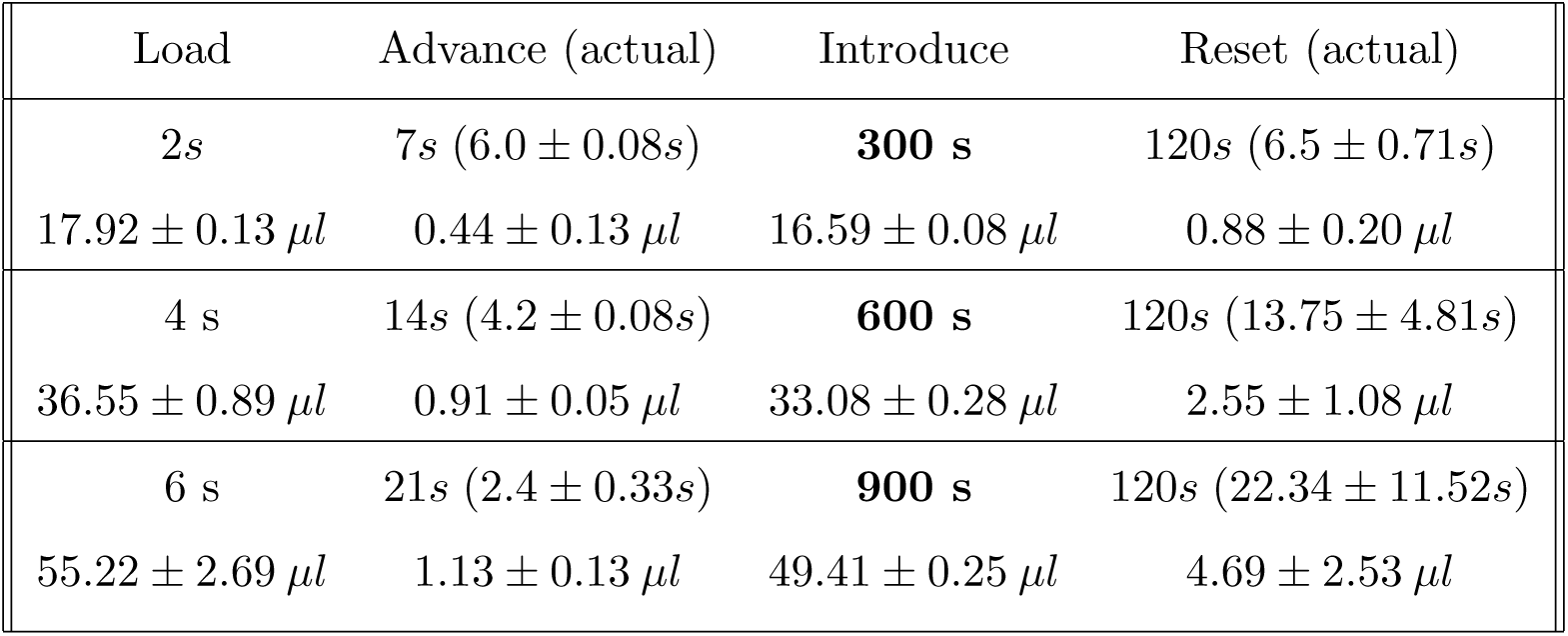
Characterization showing highly reproducible LAIR timings and volumes on our workcell. Characterized in quadruplicate, with values shown as Mean ± Standard Deviation. The protocol timings are selected proportionally based on the desired Introduce time (in bold). For the Advance stage, the actual time required for the fluid packet to reach the chip is shown in parentheses, and the volume shown is the spent volume during the Advance stage. For the Reset stage, the actual time required to clear the leftover volume is shown in parentheses, and the volume shown is the leftover volume.

| Load | Advance (actual) | Introduce | Reset (actual) |
| --- | --- | --- | --- |
| 2s<br>$17.92 \pm 0.13 \mu l$ | 7s ($6.0 \pm 0.08s$)<br>$0.44 \pm 0.13 \mu l$ | **300 s**<br>$16.59 \pm 0.08 \mu l$ | 120s ($6.5 \pm 0.71s$)<br>$0.88 \pm 0.20 \mu l$ |
| 4 s<br>$36.55 \pm 0.89 \mu l$ | 14s ($4.2 \pm 0.08s$)<br>$0.91 \pm 0.05 \mu l$ | **600 s**<br>$33.08 \pm 0.28 \mu l$ | 120s ($13.75 \pm 4.81s$)<br>$2.55 \pm 1.08 \mu l$ |
| 6 s<br>$55.22 \pm 2.69 \mu l$ | 21s ($2.4 \pm 0.33s$)<br>$1.13 \pm 0.13 \mu l$ | **900 s**<br>$49.41 \pm 0.25 \mu l$ | 120s ($22.34 \pm 11.52s$)<br>$4.69 \pm 2.53 \mu l$ |

Next, during the Advance stage (Figure 2B), the rotary valve connects one of its air inputs (i.e. selects one of two pressure controllers) to the packet of fluid resting in the output line, to advance this packet to the chip. Once the fluid reaches the chip it begins flowing out from a waste line, passing the connection to the functional region of the chip, and in this way all of the air between the fluid packet and the functional region of the chip is evacuated. Importantly, once the liquid reaches the chip, the resistance of the total fluid path increases significantly, slowing down flow (See Supplementary Video 2), and thus minimizing loss of reagent through the waste line (Supplementary section 1.2 for a more detailed explanation). As shown in Table 1, typical spent reagent volumes in the Advance stage are less than 1*µ*l, and this can be optimized further if necessary.

Subsequently, the introduction step (Figure 2C) is the critical experimental step, where we thus track the flowrate and employ pressure-based flow-control to administer a precise volume to the chip in a specified duration. Volumes administered to the chip are usually within 250 nL of each other (volumes for 5, 10, and 15 min flows, with means *±* 1 standard deviation, are shown in Table 1). In our flow-controller we use Sensirion flowrate sensors, Emerson ED02 pressure controllers, and we implement proportional integral derivative (PID) control through a Siemens Simatic S-1500 series programmable logic controller (PLC), to enable rapid and high-resolution flowrate control (Figure 2K). In this way, our software application running on a PC communicates to the PLC to operate in flow-control mode and specifies a flowrate, then the PLC containing the lower-level software manages this request in a controlled, low-latency manner, independently of the PC application (Figure 2H–I). We tuned our flow controller to operate robustly, achieving flowrate setpoints within 3s, with high stability remaining in a range of 5% of the setpoint, and across the full critical range required for our experiments (Supplementary Figure 2, see Supplementary section 1.4 for further details). Furthermore, we integrated our flowrate controller with software to perform automatic detection and response to chip clogging (discussed later).

After the introduction step is complete, the LAIR protocol ensures (through calibration of timings in Table 1) that a residual volume of reagent will remain, such that air does not enter the functional region of the chip. This volume is cleared through the waste line in the Reset stage, by flowing air in excess (usually a flow of 2 minutes), until the entire fluid path is restored to its default state of containing only air (Figure 2D, Table 1), and is therefore made independent of the next LAIR flow.

As seen in Table 1, LAIR works consistently, preventing air from entering the functional region of the chip, and producing minimal reagent loss after calibration. LAIR is implemented directly in our protocol files, through an intuitive syntax (Figure 2H). Importantly, the low resistance when the fluid path through the chip contains air allows reagents to be shuttled quickly during the Advance stage (usually taking under 6 seconds, Table 1 Advance (actual) column) which combined with the transport of fluid packets (in contrast to filling the length of tubing with fluid, necessitating a large dead volume) makes the experiment and setup independent of the distance between the reagent tubes and the chip. This allows reagents to sit far from the chip. In our workcell they are mounted to the side of a programmable microscope, and in the future we plan to scale to running chips across several different automated imaging platforms in parallel while maintaining a centralized reagent control hub. These features of the LAIR protocol, as well as alternatives to LAIR that we evaluated and decided against, are discussed in more detail in Supplementary section 1.3.

### Quake-valve control system and automated reagent mixing

Quake valves are a useful and common micromechanical element found on lab-on-chip devices (usually flexible PDMS-based) ^23^. Pressurizing a Quake valve in a control layer can collapse the device’s flow layer locally, preventing fluid from passing, and thereby providing a mechanism of active fluid routing in the flow-layer. To make the workcell compatible with multi-layer Quake-valve based microfluidics, we established an automated Quake-valve control setup similar to the Maerkl ^24, 32^, Quake ^23^, and Fordyce ^30^ labs. In particular, we adapted the well-documented and organized design from the Fordyce lab ^30^, by manufacturing arrays of water reservoirs to connect to classic arrays of solenoid valves, which makes for convenient scaling of the number of control lines (due to batched filling of all reservoirs and control lines). Solenoid valves are connected to digital output channels of our PLC.

Additionally, we integrated the ability to automate the controlled mixing of reagents (reagent mixing module) through implementing pulse-width modulation (PWM) of on-chip Quake valves using our valvecontrol system (inspired by^33^, and ^34^), which we regularly use for example to generate dilutions of a reagent during binding characterization experiments (Figure 2J, Figure 5B, E). Using this setup we have achieved concentration dilution across three orders of magnitude. Users are given control over this functionality through the protocol file using a simple syntax.

Similar to the flow-controller, the software controlling the reagent mixing module is implemented directly on the Simatic PLC, such that it can operate robustly at low latency. The backend of our desktop application simply communicates by OPCUA to the PLC-hosted server to activate the PWMmode of particular PLC channels, specifying the desired period and duty-cycle (originating from the user in the protocol file), and once the PWM step is complete, the corresponding variables in the server are flipped back to the off state. This allows for the same channel, and therefore solenoid valve and control line, to be used for both regular valve actuation and PWM. We used the PLC’s high-speed hardware pulse generators (with a minimum pulse duration of 10 microseconds) to maximize the pulse resolution and the frequency upper bound. We found that instead implementing a software solution either through the PC application or even through a cyclic program on the PLC driving the standard digital output modules results in irregular pulse waveforms or missed pulses at lower (sub 50 milliseconds) pulse durations. We characterized the outputs from the PLC using a Saleae logic analyzer (Supplementary Figure 3).

**Figure 3:**
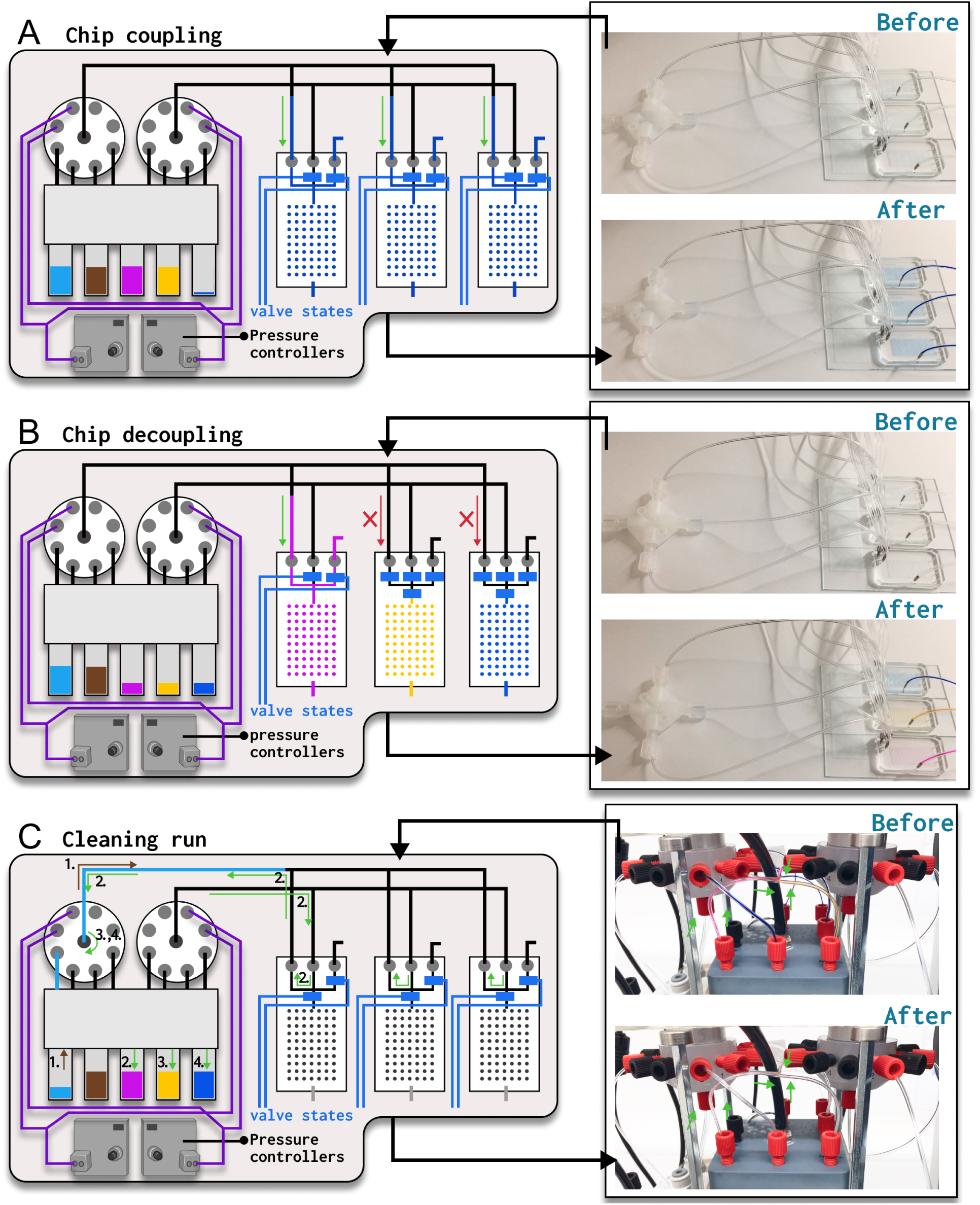
Fluid handling for lab-on-chip parallelization and setup cleaning. Caption on the next page. Figure 3: **(A-B)** Schematic depictions of how LAIR can be used to parallelize chip running, where mutliple chips are operated simultaneously, while sourcing reagents from a single copy of a reagent tube. During a single experiment, flows to parallelized chips can either be coupled **(A)** (see Supplementary Video 2), or decoupled **(B)** (see Supplementary Video 3), which is triggered dynamically depending on if the same or different reagents should flow to the chips at a given point in a protocol. **(C)** Schematic showing the fluid path used to clean rotary valve ports and reagent lines with cleaning solution at the end of every protocol. Cleaning solution is loaded into an output line between one rotary valve and the chips, and then pushed backward through the same rotary valve into its spent reagent tubes, by forming a circuit through the chips for a pressurized air line, which originates at the other rotary valve. The rotary valve rotates to direct cleaning solution to all active inlet ports, and then the rotary valve roles are reversed. A cleaning function is imported and called at the end of our protocol files.

### Microscope control

Many biological LOC experiments utilize microscopic characterization, either in brightfield or fluorescence. A fully-motorised fluorescence microscope rig is composed of numerous controllable apparatus, including lasers and lamps, configurable components along the optical train (turrets with filters, objectives, condensers), a freely programmable X,Y, and Z stage, and a good camera. Often, LOC experimenters will manually observe and image their chip under a microscope. However, this requires the experimenter to remain on standby, prevents extensive parameter-searches or over-night experiments, and limits standardization. We were interested in conducting imaging automatically, in a way that is easily programmable on a per-experiment basis, and integrated directly into our existing protocol language for cohesive experiment design. We were able to achieve this to enable users to easily program a fully motorised Nikon Eclipse Ti 2 directly from our protocol files. The user can specify steps in the experiment where the system should capture images of the chips. The software will launch NIS Elements and query the user to identify target X,Y, and Z points to image. By interpolating, our software can build a grid of points to image on each chip that is being run, based on the number of columns, rows, and chips specified. The optical configuration (magnification, fluorescence channel, exposure, etc.) can be specified in the protocol file, and parameters that are left unspecified will adopt the current default configuration in NIS. The captured images can be stored in the local file system, or a cloud storage location can be specified in the protocol file, such as an S3 bucket on AWS. Furthermore, fluidic chips sit on an Okolab glass plate that we mount on the microscope stage, which is controlled through a configurable temperature controller, such that chip temperature is precisely maintained throughout protocol execution, and to enable heated incubation steps in the experiment as specified by the user in the experiment protocol.

### Software archiecture

To manage the workcell we use a comprehensive desktop application called automancer (Python backend, Javascript frontend) that acts as a scheduler and is useful for automating laboratory processes in general. This software has been made fully open-source through an independent paper ^31^, where it is covered in detail. Control of the workcell is one demonstration of how this software can be used.

Briefly, automancer consists of a protocol editor (alternatively a Visual Studio code extension), protocol parser, executor (responsible for handling execution, communicating to the devices and to the user-interface), and an experiment dashboard for experiment management including real-time modifications to execution. To operate the workcell, we write structured text (.yaml) protocol files using a high-level, custom abstracted experiment language ^31^.

In order to execute workcell protocols, we connect the majority of our hardware to the PLC, and we write lower-level latency-critical software into the PLC directly (responsible for instance for the reagent mixing module, and the flow-controller), using Statement List (STL) and Ladder logic (LAD). We then host an OPCUA server from the PLC to expose an interface to this hardware and software, to which automancer can connect. In execution, automancer communicates desired hardware states and software parameters over OPCUA by updating variables in the server, which is read by the PLC, and the PLC communicates the actual hardware states, sensor data, and errors back, to enable tracking. The backend executes tasks asynchronously, using for instance asyncio, and natively supports parallelized execution of protocols.

Entirely new hardware can be integrated into the workcell easily, since it can be connected to a channel in the PLC, a variable is exposed on the server, and the user simply needs to update a devices’ configuration text file with the server variable name(s) and the desired parser keyword(s). Protocols in the desktop application can then immediately start using these keyword(s) as valid language for basic operation. Furthermore, automancer itself was designed with a modular architecture to foster thirdparty extensibility ^31^, and thus its use is directly in line with the previously discussed core objectives of the workcell.

For typical experiments that we run on the workcell, we fully automated image processing, noiseanalysis, parameter-inference and data visualization by developing a cloud-based pipeline through Amazon Web Services (AWS) (S3 buckets, AWS Batch and Fargate), with experiment reports containing visualizations ported into Notion for easy review. This pipeline is specific to our images, but we have included a description of the software in Supplementary section 1.7 (Supplementary Figures 4 and 5), in case it is useful as a reference.

**Figure 4:**
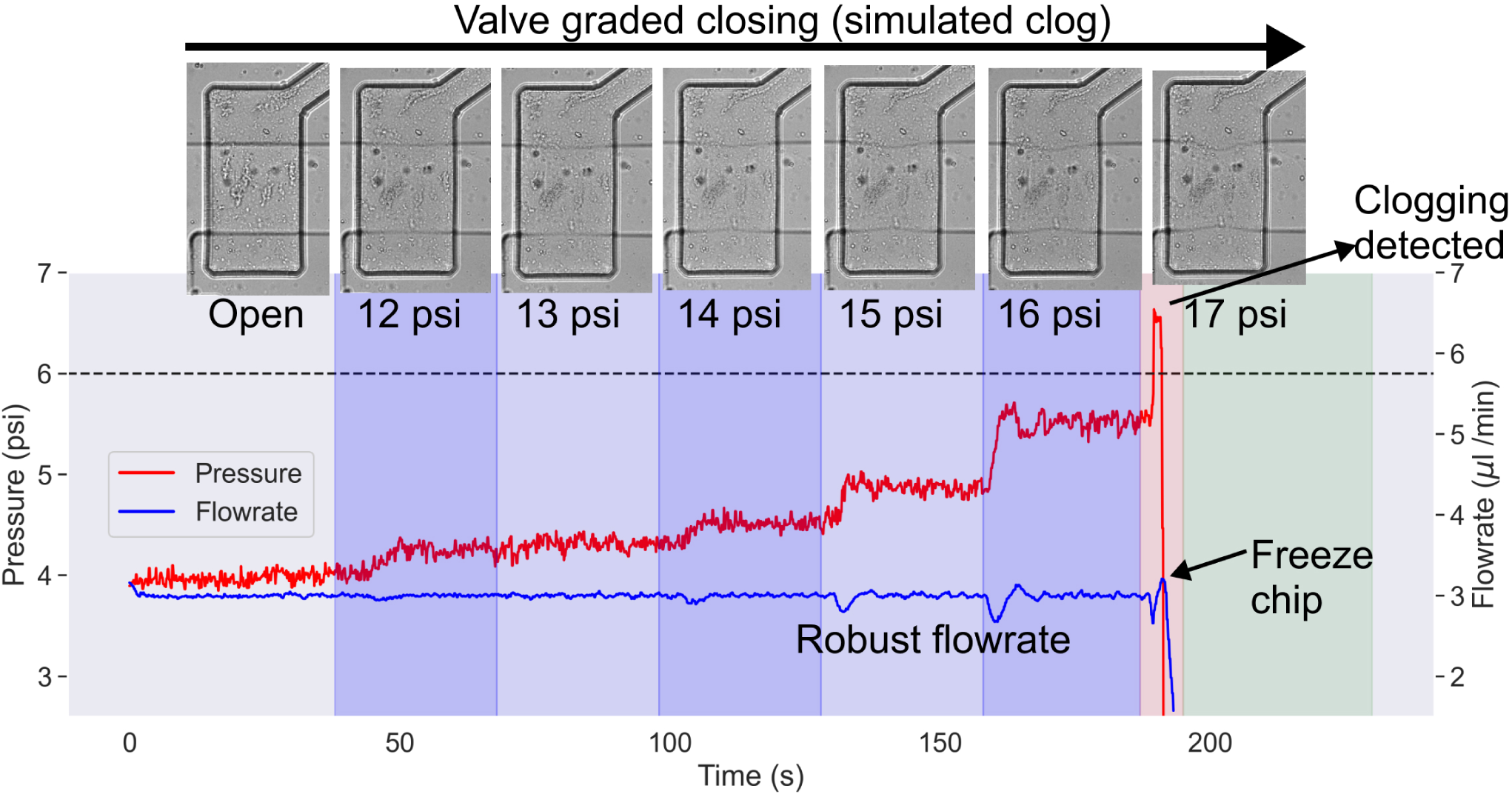
Clogging detection demonstration. We simulated a clog by progressively pressurizing a Quakevalve in the fluid path, increasing by 1 psi every 30 seconds (top). The flowrate controller increases the flowline pressure to maintain a flowrate that is robust to clogging. When clogging becomes too extreme, the clogging response is activated, and the chip is frozen into a safe state with an alert sent to the user to intervene and unclog the chip.

### Parallelization, automated cleaning

In LOC research, chips are typically run one at a time, however this limits the throughput of data collected (e.g. samples processed, sequences or biological systems characterized), and the throughput of experiment variations tested (e.g. assay variations in assay development). With automation and some plumbing it becomes possible to operate multiple chips in parallel (Figure 3A–B), whereas achieving this manually would typically require too much experimental complexity and manual loading of a prohibitively large number of reagent input tubings.

Using the workcell and LAIR, protocols can conveniently be executed on three chips in tandem (without foreseen limits on moving to higher degrees of parallelization), all the while sourcing reagents from single copies of standard labratory tubes. In our group, the goal is often to execute the same protocol across all three chips, for instance to increase the number of sequences characterized through a particular assay (up to 3072 samples for 3 chips, but in practice 1024 sequences in triplicate). In this case, we couple the LAIR flows to the chips, as demonstrated in a simplified schematic in Figure 3A and an associated Video (Supplementary Video 2).

In contrast, when developing new chips or assays, we often execute experiments in parallel that differ from one another in their protocol steps, for instance to test different versions of experiments in process development, or to characterize sequences (e.g. antibodies) against multiple different binding partners (e.g. antigens) in a single experiment. In this case, to save time we still couple the experiments where they share similarity (e.g. usually chip priming, and surface chemistry), and we decouple them where they differ (Figure 3B, Supplementary Video 2). To de-address and thus decouple chips we close their inlet Quake-valves during the Load and Introduce stages of a LAIR flow cycle (Figure 3B). Occasionally, very small volumes of liquid can enter the de-addressed tubing lines, but this volume is negligible in terms of reagent expenditure, and is promptly cleared through the waste line in the Advance and Clear stages of the LAIR cycle (which are executed as normal), as well as washed by phosphate-buffered saline (PBS) solution in the subsequent LAIR cycle.

In addition to the capacity to independently address different chips, we leverage standard multiplexing on-chip to independently address each of 32 different rows on our custom Quake-valve chips (as seen previously ^23, 35^), thus reaching 96 fully addressable flow paths in a single experiment (see Supplementary section 1.5 for further details on how we leverage multiplexing for assay development through the workcell).

Standardization should extend past experimental protocols, to cleaning procedures that are used to maintain any re-used components. To this end, we wrote a function to program automated cleaning of all re-usable components, including the flowrate sensor, both rotary valves, and the connections from the rotary valves to the pressure head (Figure 3C). This has led to improvements in the day-to-day reproducibility of flowrate measurements during protocol execution (not shown). How cleaning runs are executed on the platform is depicted in Figure 3C, and discussed in more detail in Supplementary section 1.6.

### Clogging-detection and response

A problem familiar to most Lab-on-Chip researchers is device-clogging, which is a product of working with small feature sizes and labile, aggregation-prone reagents. Clogging is an important obstacle to tackle to enable walk-away automation, since it is unpredictable, relatively frequent (in particular affecting long experiments), and typically requires user-intervention to prevent experiment failure and be resolved (except perhaps when using specific materials ^36^, or through responsive design of fluid routing (unpublished)).

We leveraged automancer’s support of clauses in protocols that cause a change in protocol execution based on a condition, in particular based on sensor input. Any protocol block can be paired with a condition (using an “expect” clause), such that if that condition becomes False, then the protocol performs some different action. In particular, we write a condition where if during regular operation, a flow controller needs to drive the pressure near to a hard maximum that is set on flow pressures (set to prevent chip delamination), then the chip is Frozen into a safe state. More precisely, the protocol reverts to the nearest protocol block that has been marked as “Stable”, where inlet valves are closed and pressures are turned off, until a user is available to intervene. The user can also be alerted, and the remaining reagent in the tubing line can even be returned to its reagent tube, by switching to a point in the protocol that initiates a fluid routing path similar to in Figure 3C.

We demonstrated this functionality in Figure 4 by conducting an experiment where we simulated a clog. We leveraged the workcell to quickly write a simple protocol where every 30s we iteratively increased the pressure applied (through a digital pressure controller) to a Quake valve to simulate a clog. Pressure on the Quake valve was increased from 12 psi to 17 psi, in 1 psi intervals (Figure 4, images on top). A flow-controller was driving reagent through the chip, past this simulated clog. The pressure plotted in Figure 4 was the flow line pressure, read from a pressure sensor that is a part of the flowrate controller. At each step of increased valve closure, the flow-controller needed to drive flow pressure higher to ensure that the flowrate is constant and thus robust to clogging. Eventually, the simulated clogging becomes too extreme (at a pressure of 17 psi on the Quake valve), and the required flow pressure threatens delamination of the flow layer (exceeding the dotted line in Figure 4). In this event, the expect clause in the protocol is triggered, and the chip reverts to a stable state where the inlet valves are closed and the pressure is turned off until the user intervenes. This combination of flowrate control paired with clogging identification and response can also be used across a wider pressure range (increasing robustness further), for devices that are plasma bonded and able to withstand higher pressures (like the chip in Figure 2G, middle).

### Workcell protocol library and versioning

The primary value of a lab is its protocols and ability to execute them. For running LOC processes through the workcell, this is made tangible as files paired with the necessary software and hardware to run them, helping with common practical issues that face labs such as skilled employees leaving the lab, learning curves for new employees, and standardization. Protocols can also then be shared more reliably with collaborating labs to make it easier to obtain the same results, provided both labs are using similar automated instruments. Furthermore, process development consists of iterating on protocols, and optimization usually comes from slowly accumulating differences over time, the history of which is usually forgotten for instance by new employees. A benefit of using a text-based experiment designer, as opposed to graphical, is that different protocol versions are naturally registered as different Git commits. Supplementary Figure 6A shows an example Git history of one of our protocols taken from our protocol registry. We associate the protocol version that was used, to the experiment data that was collected. The changes that have been made to a protocol can be viewed easily (applying a simple Git diff operation to different commits, Supplementary Figure 6B) to highlight potential causes of any differences found in experimental results.

Within automancer’s language, to expedite our writing of protocols, we have created our own library of protocol functions that are particularly useful for running lab-on-chip processes on the workcell (e.g. a function encoding a LAIR flow that accepts a reagent name and a duration as arguments, Figure 2H). We import this function library at the beginning of a protocol file, and use them to write new, readable protocols. This way of working allows us to modify the function library itself whenever a platform-general optimization is desired, or when the platform evolves and protocols need to be adjusted. These changes to the function library then propagate to all experiment files, keeping them up to date and standardized. Furthermore, compared to conventional forms of automated experiments (especially involving programmatic microscope control), where it may take weeks to establish software for a new kind of experiment or platform (often written in MATLAB or LabView), conveniently our protocols can be designed in under 30 minutes, and we expect that our experiment language can be learned in 1-2 hours by someone without prior programming expertise.

## Proof-of-concept experimentation

We demonstrate a series of 5 different automated experiments across two kinds of LOC devices that take advantage of different system features.

### Cell-free based protein expression and purification

We evaluated the ability of our platform to accomplish a challenging experiment where antibodies are first expressed directly on-chip and then purified to precise regions on the surface of the chips (Figure 5A). The experiment was designed to be run in parallel (with full coupling as in Figure 3A) across three copies of in-house designed Quake valve-based chips (relying on the working principle of Mechanically-Induced Trapping of Molecular Interactions, MITOMI^24^). The experiment was executed entirely automatically based on the LAIR flow protocol, after the initial setup by a user consisting of taking tubes from the freezer, docking them at the Hydra, and plugging in the chips. DNA encoding different synthetic antibody variants was introduced into the different “spotting chambers” of the chips during chip fabrication using an Arrayjet piezoelectric spotting robot for precise ‘x-y’ placement and volume loading.

**Figure 5:**
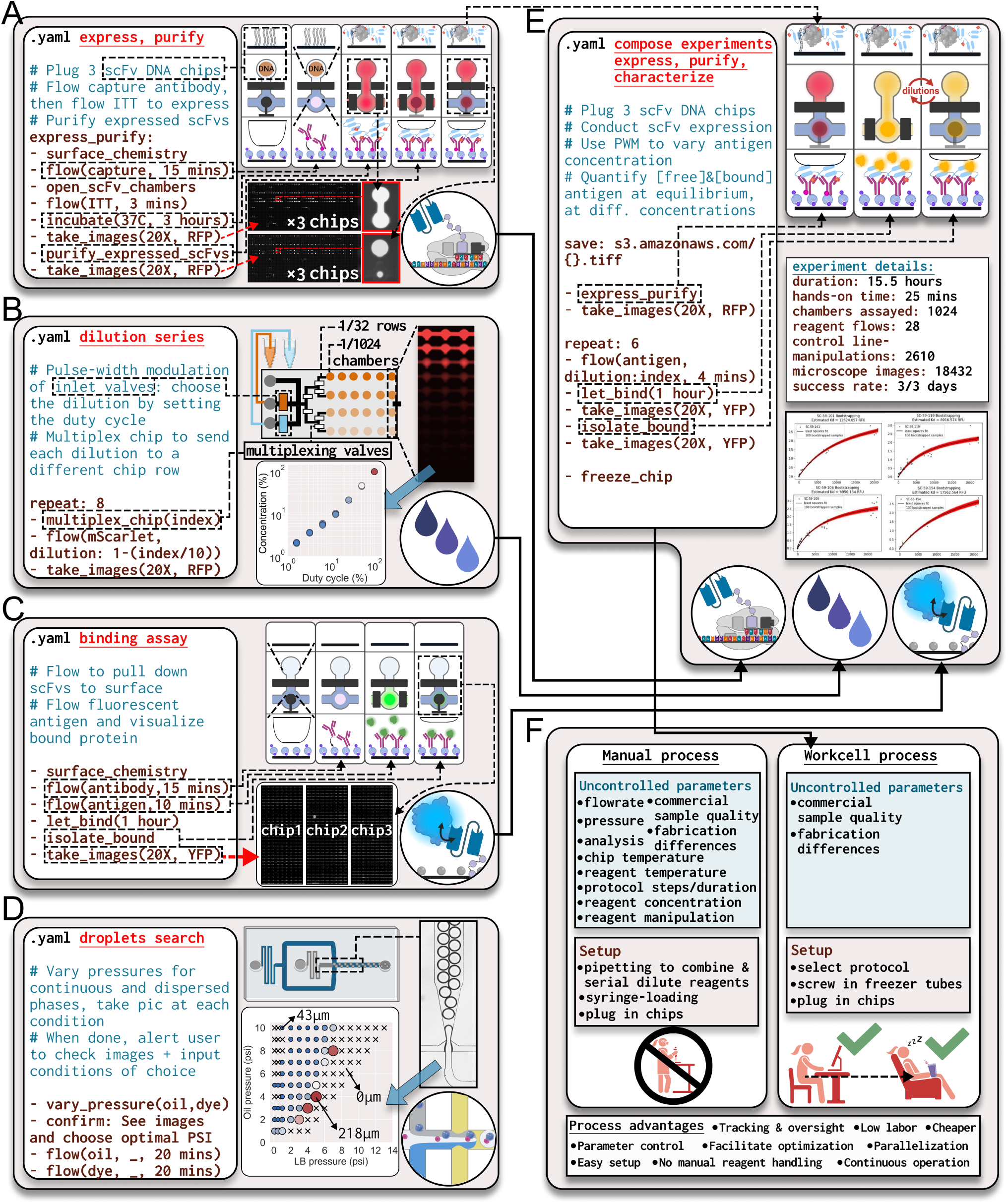
Versatile execution of a panel of different experiments. Caption on the next page. Figure 5: The panel for each experiment shows a representation of the protocol file, an experiment schematic, and results from the workcell, with arrows connecting these elements to one another to highlight their relationships. **(A)** On-chip protein expression and purification, conducted in parallel across three chips. The schematic on top shows sequential stages of the experiment, where each panel shows three views: spotting chamber (top, cross-section), full unit-cell (middle, bird’s eye view), assay chamber (bottom, cross-section). The assay chamber has a button-valve ^24^, useful to trap molecules that are pulled down to the chip surface, which is drawn at stages where it is activated. The chip was spotted with a library of scFvs to encode a different variant (DNA) in each spotting chamber. The first panel in the schematic shows the state of the unit cell after surface chemistry was conducted, followed by pulldown of capture antibody to the circular button region, cell-free expression at 37°C, and scFv purification onchip (last two panels). A full fluorescence scan of the chip is shown after incubation (top images), and after purification (bottom images), as well as a close-up image of one unit-cell at both stages. Images of the full chip are shown in larger form in Supplementary Figure 7A-B. **(B)** PWM to tune fluorescent transcription-factor protein concentration, paired with multiplexing on-chip for fluid-routing. Sample images and data show that concentration can be tuned by changing PWM duty cycle, and different concentrations can be routed to different rows of chambers. **(C)** Qualitative binding assay between IGG antibody and GFP protein, parallelized across three chips (bound protein shown in chip scans, larger form in Supplementary Figure 7C). **(D)** Droplet experiment mapping a wide range of experiment space (105 conditions) to identify accessible droplet diameters and production frequencies. A sample image is shown of droplets being generated (top right), with a plot highlighting the viable droplet space. “X” markers identify regions where droplets were not formed, while circular markers identify viable droplets. Size and color indicates drop diameter (from small and blue reflecting low diameters (min 43 *µ*m), to large and red reflecting large (max 218 *µ*m)). Subsequently, experimental parameters were selected to encapsulate single bacterial cells at 50, 120, and 190 *µ*m (Supplementary Figure 7D). **(E)** Protocols from **A, B,** and **C** were composed to execute a complex experiment involving the combined expression, purification, and quantitative binding characterization of single chain variable fragments (scFvs) against their target antigen. The schematic (top) begins from the endpoint of the schematic in A. Experiment details, and sample binding curves are shown. **(F)** Comparison between if the experiment in **E** was executed manually versus on the workcell.

As instructed in the protocol, the workcell conducted a surface chemistry on-chip, employing a “button valve” ^24^ to specifically functionalize a circular area in a detection chamber with an anti-his antibody. The workcell then opened the “spotting chambers” (Supplementary Video 4), and the dried DNA was reconstituted using a Protein synthesis Using Recombinant Elements (PURE) system (PUREfrex, Genefrontier), which is a complex and sensitive mixture of 36 recombinantly purified proteins. After it introduced the protein expression mixture, the workcell imaged the three chips across all 3072 chambers, and heated the chips to 37 degrees Celsius for 3 hours of incubation and protein expression. The button valve protecting the circular his-tag pulldown region was released 1 hour into incubation, such that expressed, his-tagged scFvs were pulled down to this circular region on the surface of the chips. The chips were imaged again under a GFP fluorescence channel (images after incubation in Figure 5A, top). The “button valves” were then pressurized, trapping pulled down scFvs, and the workcell washed the chips to purify away the unbound molecules of the complex expression mixture. The workcell imaged the chips again to characterize the purified scFvs (images after purification in Figure 5A, bottom).

### Dilution generation and multiplexed routing

We combined pulse-width modulation of inlet valves with on-chip multiplexing in order to generate a concentration gradient of fluorescent transcription factor protein across a chip (Figure 5B). Transcription factor and PBS were gated at the chip’s inlets using Quake valves. Based on the protocol file, the workcell iteratively selected a different row of the chip using multiplexing Quake valves, and then selected a different concentration of the fluorescent protein to flow to this row by ON/OFF pulsing the two reagent inlet valves such that they always had opposite states (with an XOR value of True, similar to the waveforms in Figure 2J). The duty cycle was tuned to achieve a wide range of dilutions (2 orders of magnitude shown, while we have gone to 3), where here the ON state signifies the proportion of the period where the transcription-factor inlet is open and the PBS inlet closed. The measured percent concentration (relative to the original sample) corresponded to the duty cycle set in the protocol.

### Qualitative binding experiment

To demonstrate that our platform functions as expected and is compatible with reactions found in typical biomolecular assays, we sequentially coated in parallel the surface of three chips with biotin, followed by neutravidin, and then conducted a surface-based qualitative binding reaction between antibody (antiGFP) and fluorescent protein (GFP). As seen in Figure 5C, the experiment worked as expected, and we visualized the binding of the antibody to its GFP binding partner. The experiment was fully automated through the LAIR protocol.

### Droplet microfluidics development

We were interested in establishing a novel high-throughput process for characterizing synthetic antibodies. As initial steps toward this end-goal, we decided to develop the capacity to encapsulate single bacterial cells in microfluidic droplets, from a bacterial library where different cells contain different synthetic antibody variants. We were interested in implementing this encapsulation inside of droplets spanning a wide range of diameters, due to uncertainty about the optimal droplet diameter in the downstream assay steps.

For this work, we began using new flow-focusing chips that we had not worked with before. Naturally, we exploited the workcell to automatically search the parameter-space and map the regime across which our flow-focusing chip could generate viable droplets, and to identify the range of diameters we could access, before selecting a few diameters at which to single-cell encapsulate our bacterial library. In this experiment, a protocol was written such that the workcell varied the flowrates of continuous (fluorinated oil) and dispersed (nutrient-rich Lysogeny broth (LB)) phases across 105 different conditions, capturing an image of the droplets generated at each (Figure 5D).

After analyzing the images to identify the viable droplet space and the range of accessible diameters (Figure 5D, roughly 40-220 microns), we chose settings to automate the encapsulation of single bacterial cells in droplets of 50, 120, and 190 micron diameters. We docked tubes with appropriate concentrations of bacterial cells (6’000’000 CFU/mL, 450’000 CFU/mL, and 110’000 CFU/mL respectively) each into a different port of the Hydra, such that at each diameter, based on the assumption of an ideal Poisson process, we would have roughly 26% of droplets containing 1 cell, and around 6% containing more than 1 cell. The workcell then iteratively paired the appropriate bacterial concentration with the parameters to achieve the appropriate droplet diameter. Encapsulation and characterization resulted in roughly the expected proportion of droplets containing bacterial cells for the droplets of 50 (25.2%) and 120 micron (34.9%) diameters. The 190 micron diameter droplets had a higher than expected proportion of drops with bacteria, and thus we did not move forward with this condition.

We were therefore rapidly able to encapsulate single cells in droplets across the range of diameters accessible to an unfamiliar LOC device, with minimal manual experimental effort and under 30 minutes of programming time. This highlights how the workcell enabled us to make rapid progress toward a novel application.

### Integrated expression, purification, dilution generation, quantitative binding

We further challenged our platform to evaluate if it can reliably execute long and complex experiments that are not realistic to achieve manually. Building on our past experimental results, we decided to apply our platform to integrate on a single chip the full core workflow required by a typical protein characterization lab: protein expression, protein purification, protein dilution generation, and quantitative binding characterization across multiple protein concentrations. In order to accomplish this, we wrote an integrated protocol in our software which would require 28 LAIR flows of reagents, 2610 control line manipulations of a chip containing over 3000 mechanical valves, 7 independent heated or room temperature chip-incubation steps, and 18432 high-resolution camera images to be taken across 18 scans, in brightfield and two fluorescence channels. The protocol takes 15.5 hours (930 minutes) to execute. Of this 930 minutes, only 25 minutes were manual and spent on experiment setup. We decided to run a similar experiment three days in a row, all of which executed successfully, to validate that the platform operates reliably without software or hardware bugs. Data was automatically uploaded to AWS and fed through our cloud-based image analysis pipeline (Supplementary Figures 4, and 5) to generate the binding curves in Figure 5E.

Using this experiment as an example, we listed the experimental parameters we could identify that had any possibility of varying between different runs of the same experiment to impact experimental results. We then identified which of these we had control over, if the experiment was executed manually, as compared to execution on the workcell (Figure 5F). Based on this analysis, for the experiment in Figure 5E, we reduced the uncontrolled parameters significantly from 10 identified categories in the manual case (reagent manipulation (handling), reagent concentrations, protocol steps and durations, reagent temperature, chip temperature, data analysis, pressures, flowrates, commercial sample consistency, and chip fabrication differences) to 2 identified categories remaining on the workcell (commercial sample consistency, and chip fabrication differences). More generally, compared to manual execution, LOC research conducted through the workcell benefits from parallelization, improved control over experimental conditions, parameter tracking and oversight, not requiring manual reagent handling, easier experimental setup, lower labour requirements, reduced reagent expenditure, facilitated experiment optimization, and continuous operation.

## Conclusion

Developing LOC systems usually involves the coupled design and testing of both chip architecture and biological assays on-chip, followed by optimization, and effort to increase process reproducibility and scale. Currently there is limited tooling available for chip operation. LOC devices are often utilized manually, or through DIY custom setups that offer little versatility, control over experimental conditions, or capacity for experiment design and management. Better tooling in this area could strengthen how we move from CAD file to production by parallelizing the characterization of novel device architectures and assays to expedite the discovery of properly functioning designs, by enabling researchers to easily automate optimization to move toward ideal protocols, and by improving throughput, control over experiment conditions, and tracking, to bridge the gap to production settings.

To this end, we developed a workcell that enables the design and automatic execution of experiments on LOC devices. We were particularly interested in satisfying the operation of more complex systems relying on active fluid routing, where abstraction of experiment design and comprehensive automation seemed essential. We focused on developing a starter pack of modules that together support a range of different functionalities, including reagent cooling and transport (by LAIR), reagent mixing and dilutions, flowrate and pressure control, clogging-detection, on-chip valve control, automated microscopy, chip-temperature control, and automated setup cleaning, all controlled seamlessly from the same easy-to-write protocol file. Leveraging this functionality, we demonstrated that our platform is versatile, by performing considerably different applications across both Quake-valve and droplet microfluidic systems.

We emphasized how the workcell supports parallelized and addressable chip-operation, which can be used to test a greater number of chip or experiment designs in parallel, or simply to increase process throughput. We showed that the workcell can be rapidly programmed to run parameter-search experiments for device characterization and selection of optimal operating parameters. Additionally, with only 25 minutes of initial user setup time, the workcell implemented a highly complex, 15.5 hour long experiment successfully, and this was accomplished three experimental days in a row. Use of the workcell enabled us to dramatically reduce the number of uncontrolled parameters in this experiment, and to use comprehensive tracking and protocol versioning to connect changes in results back to experimental differences.

Despite the demonstrated versatility of our platform and significant similarity in requirements across different LOC applications, the workcell will not work out of the box for certain applications (in particular, where different actuators are preferred for fluid routing, or detectors for characterization), and furthermore other labs interested in establishing their own workcell may have additional features that they wish to integrate either as hardware or software modules. We thus believe that an attempt at a general chip-operating platform would be best tackled as an open-source effort where a focus on modular hardware and software architectures affords extensibility. The LOC workcell was designed with this in mind, in contrast to custom setups that are difficult to internalize and reprogram.

Due to the simplicity of automancer’s experiment language, other labs can easily design their own protocols, including researchers having no prior programming expertise. More involved platform extension is also possible. The desktop software is largely composed of different interacting modules ^31^, and new hardware can be connected to the PLC, which has its own scalable architecture that can be matched to users’ needs. By updating a simple configuration file the parser will understand new keywords of interest, such that experiments can immediately be designed to control newly integrated hardware. We built upon many past contributions in the LOC community ^23, 24, 30, 33, 37, 38^, and we aim to support other labs to use and contribute to this project.

In addition to collaboration benefiting the development and refinement of such a platform, we also believe that automation is a good solution to improve collaboration between labs and to increase synergy in biological research. In the development of general automation platforms, an emphasis on accessibility and affordability may be required for adoption across a high proportion of biology labs, and in the future this adoption may be the necessary kick to improve repeatability and re-usability of research across labs, which at present the biological research community appears to lack^39^ compared to for instance software development communities. The combination of automation and inputs standardization could one day land us somewhere between the status quo and a requirements.txt file.

Through the workcell, both the complexity and the required time to setup chips requiring multiple reagents is reduced significantly, and we expect that the platform can be copied and parallelized while still being operated by a single employee, keeping operating costs low. Furthermore, by building the workcell from scratch using components sourced directly from manufacturers where possible, we minimized the fixed cost of the full setup. In fact, the full platform cost that we estimate is less than the typical price charged by a CRO to a commercial partner in industry for a drug discovery project, creating an opportunity for rapid self-funded copying of the workcell in a commercial setting.

In addition to improvements in chip-operation, LOC development will benefit from advances in CAD tools, as well as microfabrication approaches that offer rapid prototyping, increased flexibility, and reproducibility in the manufacture of complex multi-layer devices (e.g. avoiding alignments). For instance, large language models (LLMs)^40, 41^ and other machine learning tools ^42^ show promise for streamlining the design of complex chip architectures and for generating many designs in parallel for high-throughput prototyping, while approaches like 2-photon lithography-based 3D printing ^43^ may offer increased flexibility, for instance for design of valves^44, 45^ and fluid routing in 3 dimensions. At present, the degree to which LOC modules can be composed into systems, and how much empirical testing will ultimately be required in design, remains an open question. Improved platforms like the LOC workcell for general and parallelized chip-operation will play a role in enabling the scalable characterization and optimization of individual fluidic modules and their integrations, to prevent bottlenecking as CAD^46^ and fabrication tools ^47^ are made increasingly compatible with device prototyping. In this way, the LOC researcher will be well-positioned to understand system composability, solve associated underlying challenges, and approach a future where it becomes more common to implement a necessary laboratory workflow by integrating a set of on-chip modules. Despite being a bit slow out of the gates, progress in foundational technologies has made it an exciting time to work with lab-on-chip systems, and we believe LOCs will play an important role in the labs of the future.

## Author contributions

A.S. conceived of, designed the workcell, and directed the work. A.S. and C.N. assembled and characterized the system, designed the animation, developed the workcell CAD files and manufacturing thereof. G.B. performed all experiments using the platform, and conducted wafer and chip-fabrication. A.S. developed the chip designs and CAD files for photolithography. A.S. and G.B. wrote experiment protocols, optimized system performance and designed the automated cleaning circuit. C.N. programmed the programmable logic controller and the animation. S.L. wrote the core software application including the user-interface, and L.H. wrote the image analysis pipeline with support from A.S. A.S. wrote the paper with review from all authors.

## Code availability

Code is available in the following GitHub repository (https://github.com/eukaryoting/LOC-workcell).

## Supporting information

CAD animation of the workcell

LAIR parallelization (coupling)

LAIR parallelization (decoupling)

Automated spotting chamber filling

## Acknowledgments

A.S. and C.N. would like to thank Andreas Rohrbach from Siemens automation for enabling rapid procurement and support. A.S. would like to thank Lars Kraus and Stéphane Klich from Emerson automation.

## References

[1] P. Wang, L. Robert, J. Pelletier, W. L. Dang, F. Taddei, A. Wright, and S. Jun, “Robust growth of escherichia coli,” Current biology, vol. 20, no. 12, pp. 1099–1103, 2010.

[2] A. Groisman, C. Lobo, H. Cho, J. K. Campbell, Y. S. Dufour, A. M. Stevens, and A. Levchenko, “A microfluidic chemostat for experiments with bacterial and yeast cells,” Nature methods, vol. 2, no. 9, pp. 685–689, 2005.

[3] A. S. Rajkumar, N. Dénervaud, and S. J. Maerkl, “Mapping the fine structure of a eukaryotic promoter input-output function,” Nature genetics, vol. 45, no. 10, pp. 1207–1215, 2013.

[4] H. Ming Yip, S. Cheng, E. J. Olson, M. Crone, and S. J. Maerkl, “Perfect adaptation achieved by transport limitations governs the inorganic phosphate response in s. cerevisiae,” Proceedings of the National Academy of Sciences, vol. 120, no. 2, p. e2212151120, 2023.

[5] M. Margulies, M. Egholm, W. E. Altman, S. Attiya, J. S. Bader, L. A. Bemben, J. Berka, M. S. Braverman, Y.-J. Chen, Z. Chen, et al., “Genome sequencing in microfabricated high-density picolitre reactors,” Nature, vol. 437, no. 7057, pp. 376–380, 2005.

[6] M.-J. R. Shen, R. C. Kain, K. M. Kuhn, A. H. Talasaz, and A. Jamshidi, “Integrated sequencing apparatuses and methods of use,” Jan. 28 2014. US Patent 8,637,242.

[7] S. N. Bhatia and D. E. Ingber, “Microfluidic organs-on-chips,” Nature biotechnology, vol. 32, no. 8, pp. 760–772, 2014.

[8] S. Haeberle and R. Zengerle, “Microfluidic platforms for lab-on-a-chip applications,” Lab on a Chip, vol. 7, no. 9, pp. 1094–1110, 2007.

[9] N. Convery and N. Gadegaard, “30 years of microfluidics,” Micro and Nano Engineering, vol. 2, pp. 76–91, 2019.

[10] G. M. Whitesides, “The origins and the future of microfluidics,” nature, vol. 442, no. 7101, pp. 368– 373, 2006.

[11] H. H. Caicedo and S. T. Brady, “Microfluidics: the challenge is to bridge the gap instead of looking for a ‘killer app’,” Trends in Biotechnology, vol. 34, no. 1, pp. 1–3, 2016.

[12] D. S. Kong, T. A. Thorsen, J. Babb, S. T. Wick, J. J. Gam, R. Weiss, and P. A. Carr, “Open-source, community-driven microfluidics with metafluidics,” Nature biotechnology, vol. 35, no. 6, pp. 523–529, 2017.

[13] C. M. Klapperich, “Microfluidic diagnostics: time for industry standards,” Expert review of medical devices, vol. 6, no. 3, pp. 211–213, 2009.

[14] H. Van Heeren, “Standards for connecting microfluidic devices?,” Lab on a Chip, vol. 12, no. 6, pp. 1022–1025, 2012.

[15] D. R. Reyes, H. van Heeren, S. Guha, L. Herbertson, A. P. Tzannis, J. Ducŕee, H. Bissig, and H. Becker, “Accelerating innovation and commercialization through standardization of microfluidic-based medical devices,” Lab on a Chip, vol. 21, no. 1, pp. 9–21, 2021.

[16] E. K. Sackmann, A. L. Fulton, and D. J. Beebe, “The present and future role of microfluidics in biomedical research,” Nature, vol. 507, no. 7491, pp. 181–189, 2014.

[17] A. Y. Fu, C. Spence, A. Scherer, F. H. Arnold, and S. R. Quake, “A microfabricated fluorescence-activated cell sorter,” Nature biotechnology, vol. 17, no. 11, pp. 1109–1111, 1999.

[18] D. C. Duffy, J. C. McDonald, O. J. Schueller, and G. M. Whitesides, “Rapid prototyping of microfluidic systems in poly (dimethylsiloxane),” Analytical chemistry, vol. 70, no. 23, pp. 4974–4984, 1998.

[19] D. Wu, J. Qin, and B. Lin, “Electrophoretic separations on microfluidic chips,” Journal of Chromatography A, vol. 1184, no. 1-2, pp. 542–559, 2008.

[20] A. J. Mach, J. H. Kim, A. Arshi, S. C. Hur, and D. Di Carlo, “Automated cellular sample preparation using a centrifuge-on-a-chip,” Lab on a Chip, vol. 11, no. 17, pp. 2827–2834, 2011.

[21] C. D. Ahrberg, A. Manz, and B. G. Chung, “Polymerase chain reaction in microfluidic devices,” Lab on a Chip, vol. 16, no. 20, pp. 3866–3884, 2016.

[22] J. W. Hong, V. Studer, G. Hang, W. F. Anderson, and S. R. Quake, “A nanoliter-scale nucleic acid processor with parallel architecture,” Nature biotechnology, vol. 22, no. 4, pp. 435–439, 2004.

[23] T. Thorsen, S. J. Maerkl, and S. R. Quake, “Microfluidic large-scale integration,” Science, vol. 298, no. 5593, pp. 580–584, 2002.

[24] S. J. Maerkl and S. R. Quake, “A systems approach to measuring the binding energy landscapes of transcription factors,” Science, vol. 315, pp. 233–237, Jan. 2007.

[25] C. Markin, D. Mokhtari, F. Sunden, M. Appel, E. Akiva, S. Longwell, C. Sabatti, D. Herschlag, and P. Fordyce, “Revealing enzyme functional architecture via high-throughput microfluidic enzyme kinetics,” Science, vol. 373, no. 6553, p. eabf8761, 2021.

[26] E. Z. Macosko, A. Basu, R. Satija, J. Nemesh, K. Shekhar, M. Goldman, I. Tirosh, A. R. Bialas, N. Kamitaki, E. M. Martersteck, et al., “Highly parallel genome-wide expression profiling of individual cells using nanoliter droplets,” Cell, vol. 161, no. 5, pp. 1202–1214, 2015.

[27] R. Zilionis, J. Nainys, A. Veres, V. Savova, D. Zemmour, A. M. Klein, and L. Mazutis, “Single-cell barcoding and sequencing using droplet microfluidics,” Nature protocols, vol. 12, no. 1, pp. 44–73, 2017.

[28] F. Lan, B. Demaree, N. Ahmed, and A. R. Abate, “Single-cell genome sequencing at ultra-highthroughput with microfluidic droplet barcoding,” Nature biotechnology, vol. 35, no. 7, pp. 640–646, 2017.

[29] R. Gaa, E. Menang-Ndi, S. Pratapa, C. Nguyen, S. Kumar, and A. Doerner, “Versatile and rapid microfluidics-assisted antibody discovery,” in MAbs, vol. 13, p. 1978130, Taylor & Francis, 2021.

[30] K. Brower, R. R. Puccinelli, C. J. Markin, T. C. Shimko, S. A. Longwell, B. Cruz, R. Gomez-Sjoberg, and P. M. Fordyce, “An open-source, programmable pneumatic setup for operation and automated control of single-and multi-layer microfluidic devices,” HardwareX, vol. 3, pp. 117–134, 2018.

[31] S. Lietar, G. Bunke, G. Michielin, A. Shahein*, and S. J. Maerkl*, “Design and automatic control of laboratory experiments through automancer,” unpublished, 2023.

[32] M. Geertz, D. Shore, and S. J. Maerkl, “Massively parallel measurements of molecular interaction kinetics on a microfluidic platform,” Proceedings of the National Academy of Sciences, vol. 109, no. 41, pp. 16540–16545, 2012.

[33] K. Woodruff and S. J. Maerkl, “Microfluidic module for real-time generation of complex multi-molecule temporal concentration profiles,” Analytical chemistry, vol. 90, no. 1, pp. 696–701, 2018.

[34] Z. Swank and S. J. Maerkl, “Cfpu: a cell-free processing unit for high-throughput, automated in vitro circuit characterization in steady-state conditions,” BioDesign Research, vol. 2021, 2021.

[35] I. E. Araci and S. R. Quake, “Microfluidic very large scale integration (mvlsi) with integrated micromechanical valves,” Lab on a Chip, vol. 12, no. 16, pp. 2803–2806, 2012.

[36] C. Murray, D. McCoul, E. Sollier, T. Ruggiero, X. Niu, Q. Pei, and D. D. Carlo, “Electro-adaptive microfluidics for active tuning of channel geometry using polymer actuators,” Microfluidics and nanofluidics, vol. 14, pp. 345–358, 2013.

[37] D. Irimia, D. A. Geba, and M. Toner, “Universal microfluidic gradient generator,” Analytical chemistry, vol. 78, no. 10, pp. 3472–3477, 2006.

[38] Y. Xia and G. M. Whitesides, “Soft lithography,” Annual review of materials science, vol. 28, no. 1, pp. 153–184, 1998.

[39] C. G. Begley and L. M. Ellis, “Raise standards for preclinical cancer research,” Nature, vol. 483, no. 7391, pp. 531–533, 2012.

[40] Y. Ganin, S. Bartunov, Y. Li, E. Keller, and S. Saliceti, “Computer-aided design as language,” Advances in Neural Information Processing Systems, vol. 34, pp. 5885–5897, 2021.

[41] T. Galanos, A. Liapis, and G. N. Yannakakis, “Architext: Language-driven generative architecture design,” arXiv preprint arXiv:2303.07519, 2023.

[42] D. McIntyre, A. Lashkaripour, P. Fordyce, and D. Densmore, “Machine learning for microfluidic design and control,” Lab on a Chip, vol. 22, no. 16, pp. 2925–2937, 2022.

[43] P. F. van Altena and A. Accardo, “Micro 3d printing elastomeric ip-pdms using two-photon polymerisation: A comparative analysis of mechanical and feature resolution properties,” Polymers, vol. 15, no. 8, p. 1816, 2023.

[44] S. J. Keating, M. I. Gariboldi, W. G. Patrick, S. Sharma, D. S. Kong, and N. Oxman, “3d printed multimaterial microfluidic valve,” PloS one, vol. 11, no. 8, p. e0160624, 2016.

[45] H. Gong, A. T. Woolley, and G. P. Nordin, “High density 3d printed microfluidic valves, pumps, and multiplexers,” Lab on a Chip, vol. 16, no. 13, pp. 2450–2458, 2016.

[46] S. H. Hong, H. Yang, and Y. Wang, “Inverse design of microfluidic concentration gradient generator using deep learning and physics-based component model,” Microfluidics and Nanofluidics, vol. 24, pp. 1–20, 2020.

[47] S. Razavi Bazaz, O. Rouhi, M. A. Raoufi, F. Ejeian, M. Asadnia, D. Jin, and M. Ebrahimi Warkiani, “3d printing of inertial microfluidic devices,” Scientific reports, vol. 10, no. 1, p. 5929, 2020.

