## Supplementary material for "A workcell 1.0 for programmable and controlled operation of multiple fluidic chips in parallel"

[2]École Polytechnique Fédérale de Lausanne (EPFL), Lausanne, Switzerland

April 16th, 2023

*These authors contributed equally to this work

### Supplementary figures and tables

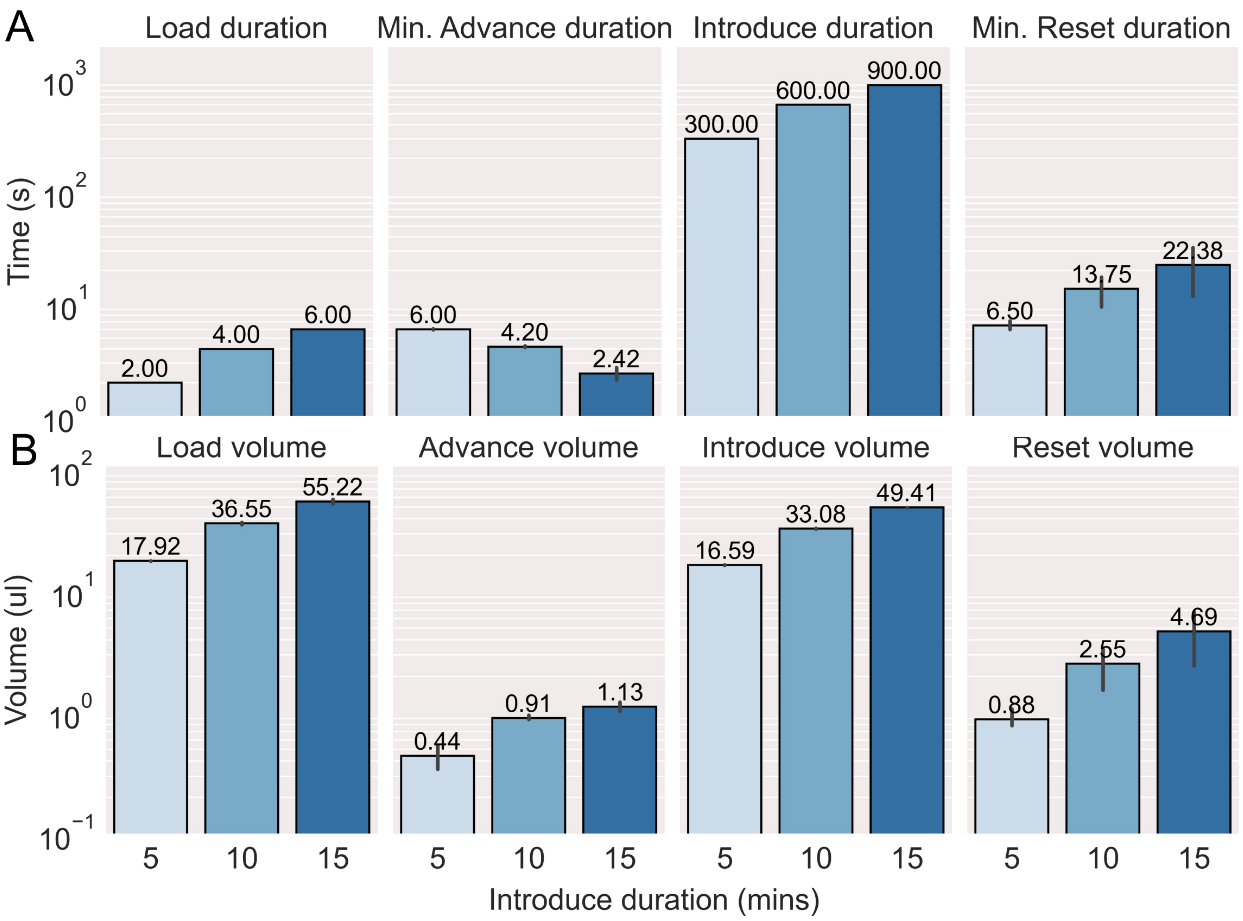


Figure 1: This plot is a visual representation of the data in Table 1; characterization showing timings and volumes for the Load Advance Introduce Reset (LAIR) flow protocol executed on the workcell. Values are means with errorbars representing 95% confidence intervals.

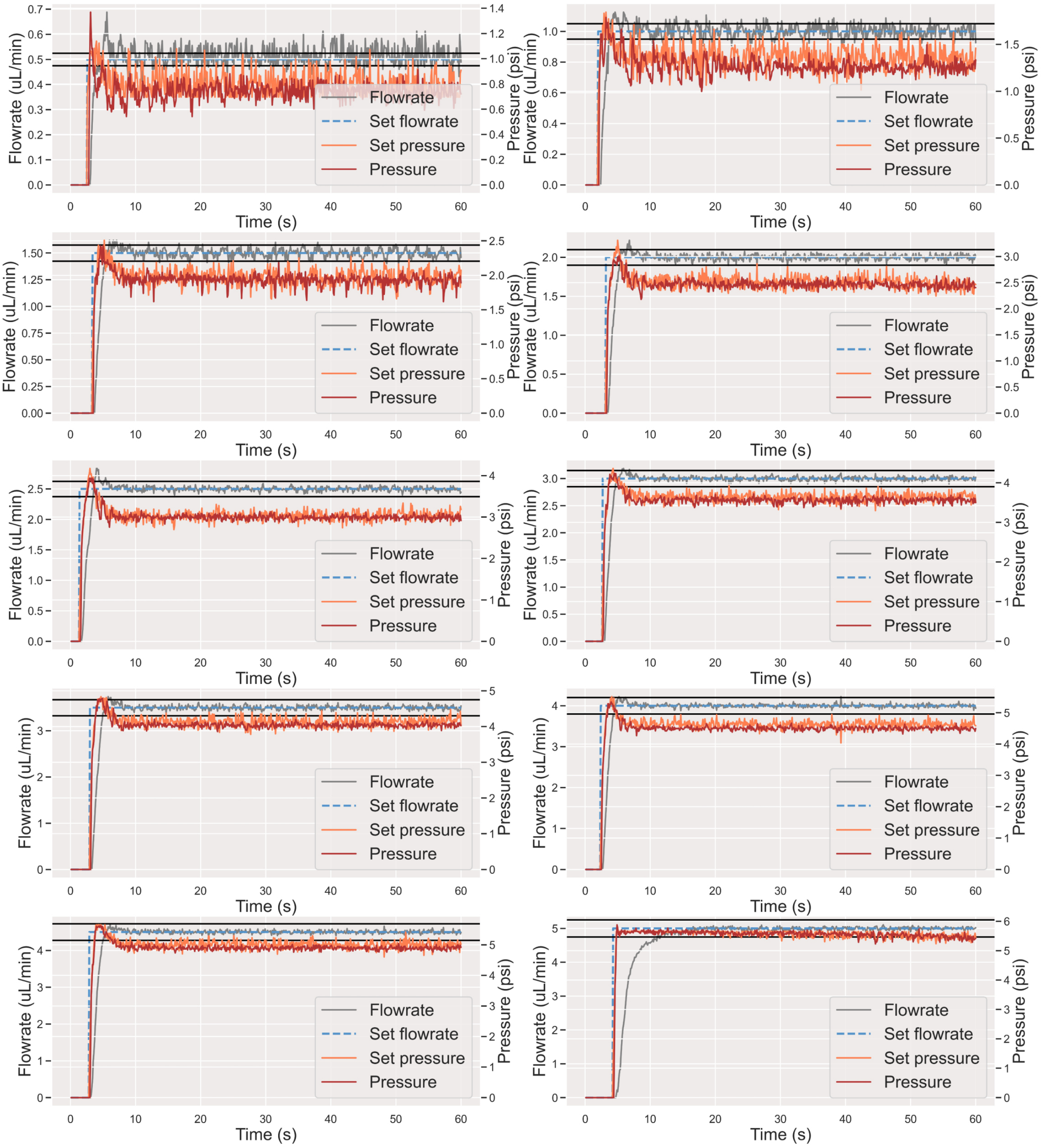


Figure 2: Characterization showing operation of our flowrate controller across a range of flowrate setpoints ($0.5\mu$l/min to $5\mu$l/min). Black horizontal lines show the $\pm5\%$ region arouund the setpoint. Setpoints are achieved rapidly, with minimal error.

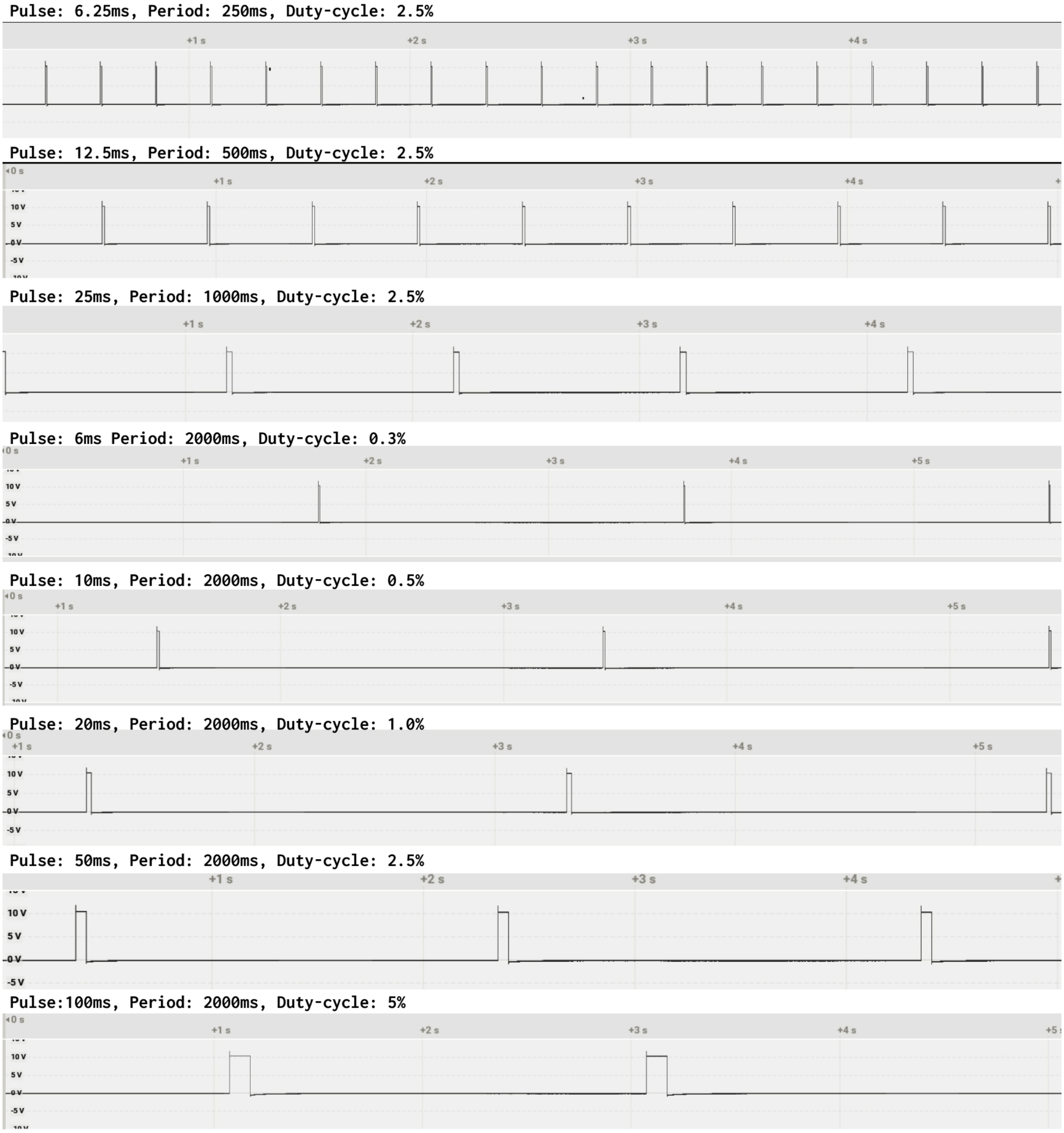


Figure 3: Characterization of pulse-width modulation outputs from the programmable logic controller, using a Saleae logic analyzer. Voltage pulses are consistent, characteristic square waves, which was not the case when using standard digital output channels from the PLC, or when attempting to drive PWM through desktop software.

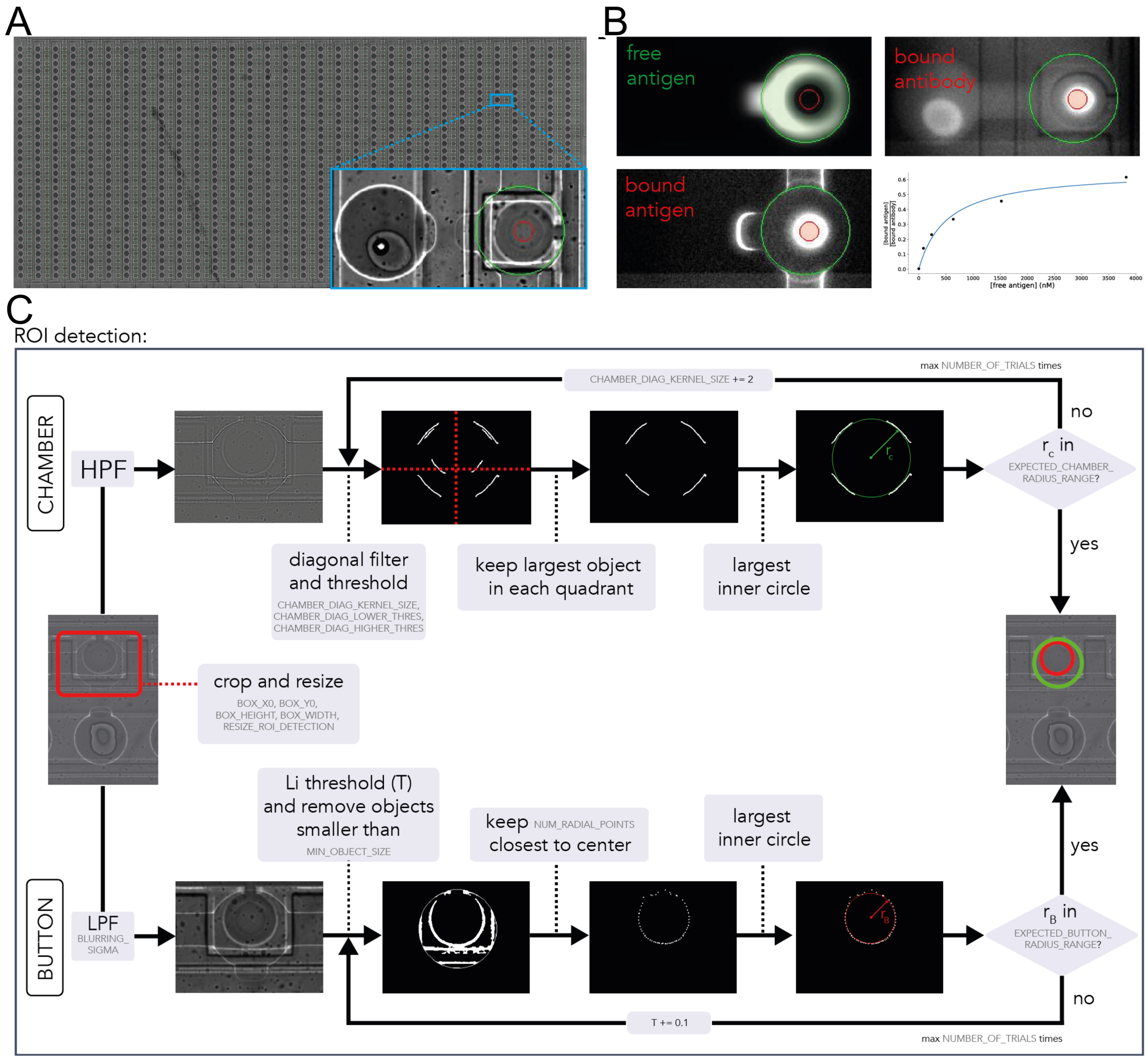


Figure 4: Overview of the ROI-detection method. **(A)** Stitched brightfield scan of a full chip. The brightfield images provide high enough contrast to detect the chamber and button ROIs (see inset). The results of the ROI detection are indicated as red and green circles for the button and chamber ROIs, respectively. **(B)** Representative fluorescent images of free antigen (taken before the antigen is washed away, but with the button down in contrast to[1], to improve consistency of the valve's position), bound antigen (after wash), and bound antibody. Regions where the fluorescence is quantified are shaded. The binding curve is the result of these scans taken at varying antigen concentrations. **(C)** Graphical representation of the ROI-detection algorithm as explained in the text.

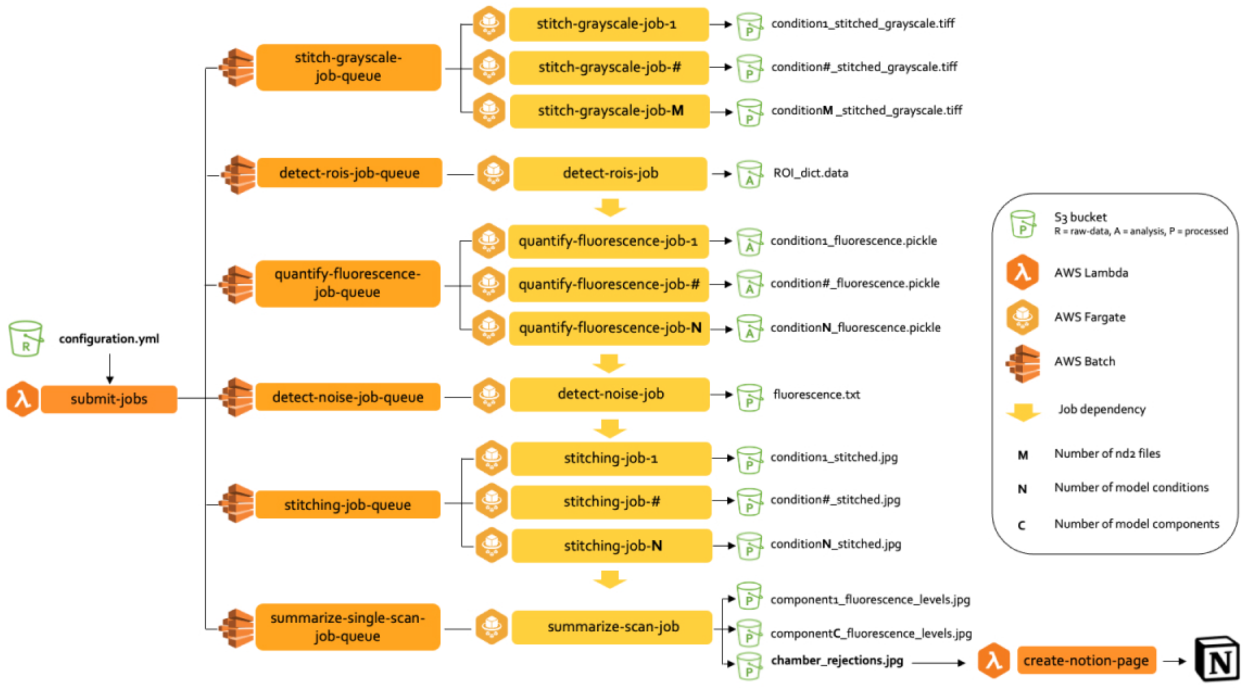


Figure 5: Implementation of ROI-detection and quantification on Amazon Web Services (AWS), using S3, Batch and Fargate.

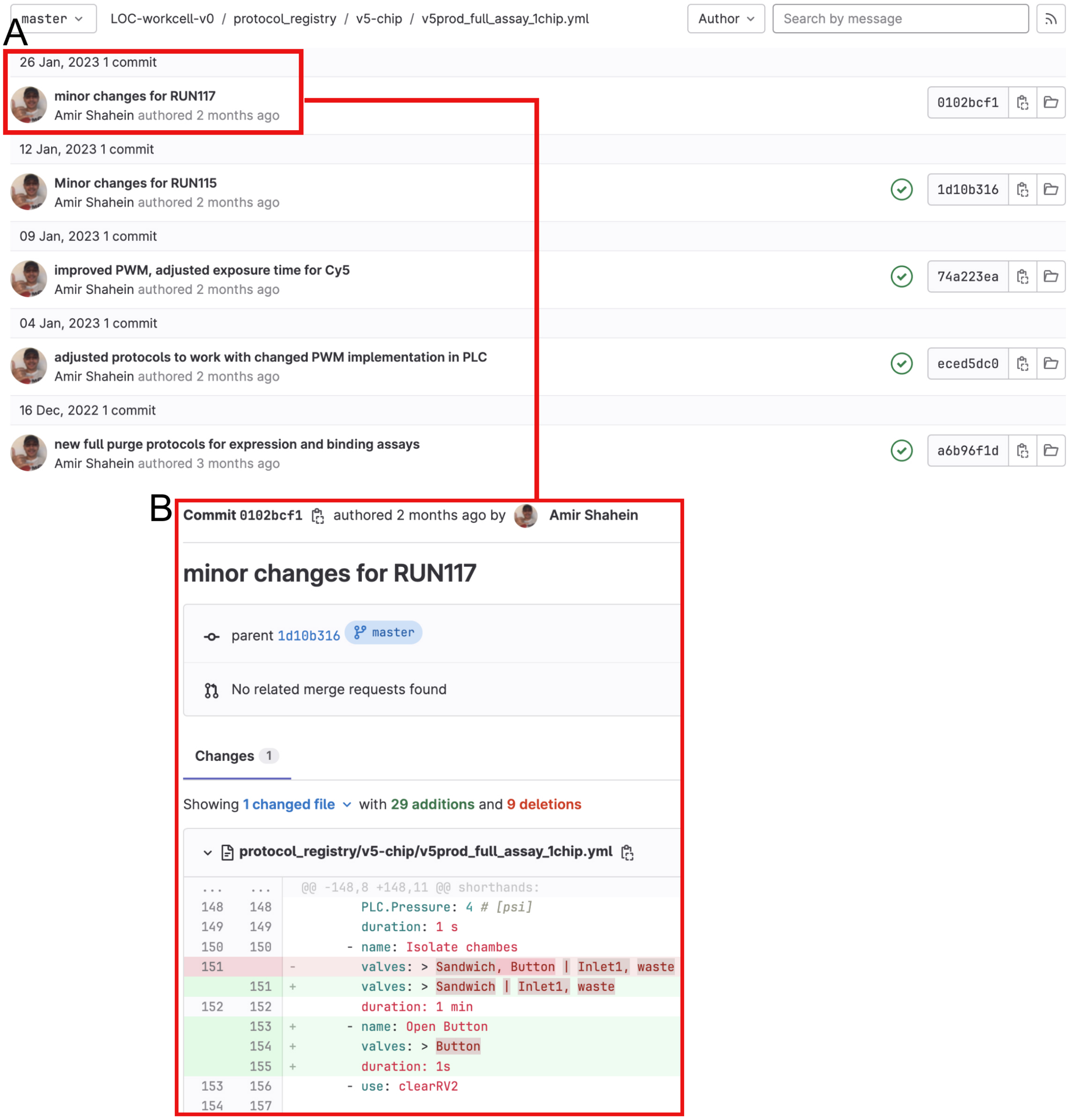


Figure 6: Example showing the Git-based versioning history of one of the automated protocols from our protocol registry (top). **(A)** A series of protocol changes are recorded as commits (versions). **(B)** Each of these commits can be opened to see the precise changes to the protocol that were made (git diff). Associating protocol versions to the experimental data, allows for easy identification of impacts on experimental results.

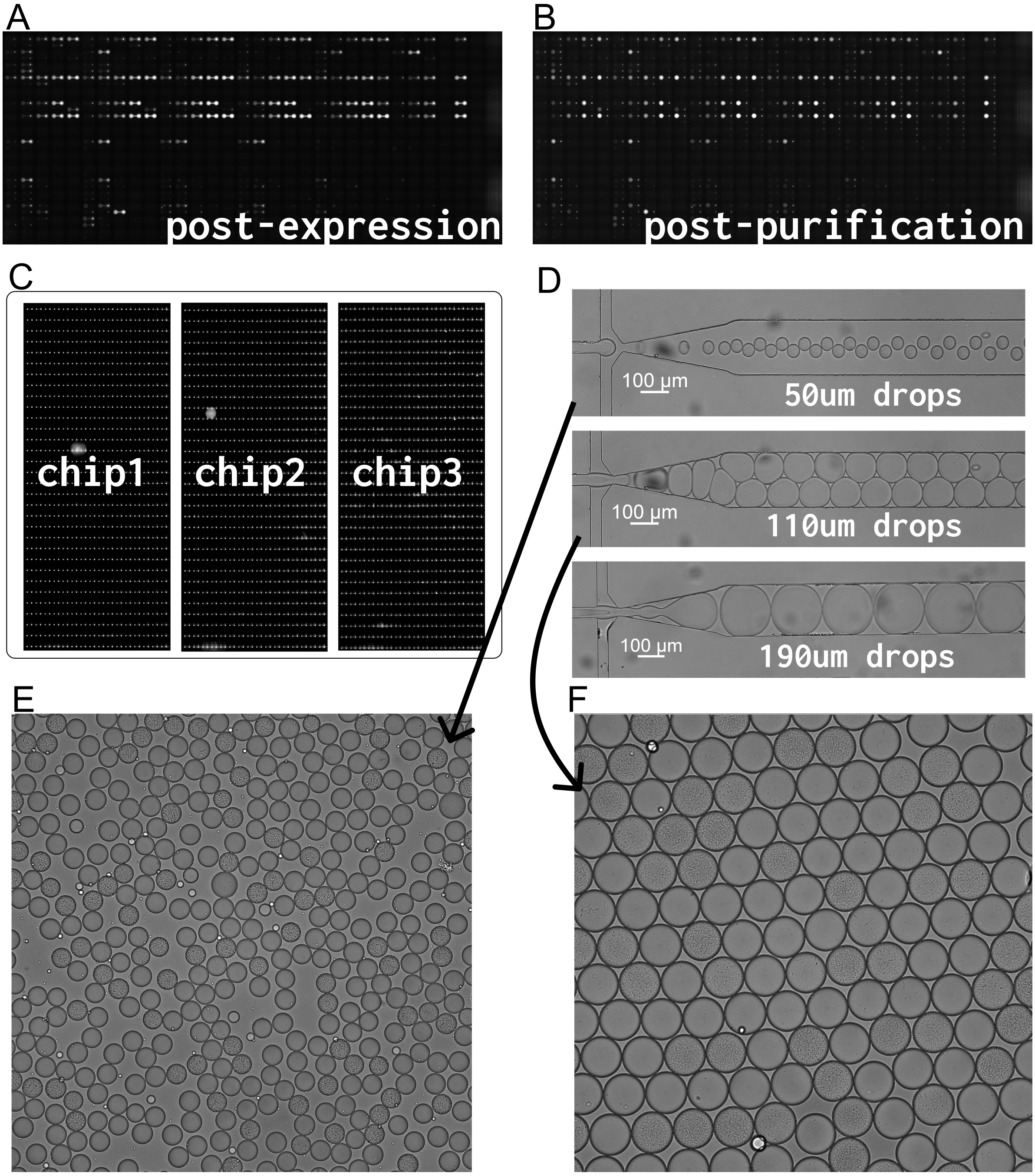


Figure 7: Raw image data corresponding to Figure 5. Scan of a chip from Main Figure 5A after protein expression **(A)**, and then after purification **(B)**. **(C)** Chips from Main Figure 5C. **(D)** Single-cell encapsulation of chips at 50$\mu$m, 110$\mu$m, and 190$\mu$m from Main Figure 5D. **(E)** 50$\mu$ droplets after bacterial growth, showing 87/345 droplets with bacteria. **(F)** 110$\mu$ droplets after bacterial growth, showing 52/149 droplets with bacteria.

### 1 Platform design

#### 1.1 Hydra fluid control system

The hydra fluid control system enables the user to simply take reagent aliquots from a freezer, and screw them directly into the assembly's pressure head. A seal is formed as the tube is screwed into the thread, since the bottom of the pressure head will compress the reagent tube's built-in O-ring against its plastic. One position in the pressure head is reserved for a sensitive resistance-based temperature sensor sitting permanently in a dummy tube containing water instead of reagent. The loaded reagent tubes mate with a custom manufactured aluminum cooling block, where they sit enclosed for the remainder of the experiment. After Computer Numerical Control (CNC) milling, PDMS was permanently casted all around the aluminum block for insulation. The aluminum block sits on a 2 by 2 array of Peltier elements, which we joined with conductive heat paste to the aluminum on top, and on the bottom to a metallic water head from a GPU cooling system. Peltier elements with suitable performance were selected based on the anticipated heat load and desired temperature difference. Liquid coolant is pumped through the water head to drain heat from the hot side of the Peltier array, and this heat is then dissipated through three fans. The water head, the pump, and each fan has a circle of addressable LEDs. We programmed the water head, pump, and one fan to report the temperature of the reagent tubes (based on a color gradient from pink to light blue), and we programmed the other two fans to report the progress of the current step in the protocol, and the experiment's total progress (by proportional filling of white circles).

On the other side of the pressure head, a seal is formed against output tubing, which connects from each reagent tube into the inputs of one of two rotary distributor valves. Together the rotary valves accept 24 inputs and deliver 2 outputs. Maintaining integrity of reagents was important, and so our design ensures that a reagent does not come into contact with any internal surfaces of the pressure head during ejection, apart from a standard biocompatible tubing of choice such as PTFE or tygon. Reagents that may need to be mixed together in the experiment connect into different rotary valves. We also mounted additional larger volume 3D-printed reservoirs (not shown in the animation) that connect to the rotary valves, containing larger supplies of solutions that are not precious and that don't need to be cooled, for instance non-biological cleaning solutions such as ethanol and PBS (explained in more detail in the automated cleaning section), or oil and water for very long experiemnts in droplet microfluidics. Furthermore, two pressurized air lines connect as input to each rotary valve, controlled by two pressure controllers (each with two outputs). Air pressure lines travelling to the pressure head, to the rotary valves, and to the cleaning solution reservoirs are all switchable through a bank of digital solenoid valves, which are downstream of one of the pressure controllers. The two pressure controllers enable separate control of flow for liquids flowing through different rotary valves, to allow different reagents that are flowing on-chip simultaneously to have different flowrates, which is useful for instance in droplet microfluidics applications.

Through the hydra fluid control system combined with the reagent mixing module, we bypass the need for manual reagent pipetting and reagent loading entirely. At this point, chip setup comprises plugging in either just one or two empty tubing lines into each chip (connecting to rotary valve outputs), as well as control lines filled with water, for which differences in manual implementation likely do not impact the experiment. Feeding all reagents through one or two inlet lines allows us to cut down the number of reagent inlet lines considerably, and therefore also the corresponding control lines that gate the inlets. Together with bypassing syringe reagent loading, we decreased the setup time for our standard Quake-valve chip to a point that 3 chips can now be connected in roughly the time it previously took for a single chip.

#### 1.2 Advance stage: increase in resistance

Once liquid reaches the chip, the resistance of the total fluid path increases significantly, slowing down flow (Supplementary Video 2), and thus minimizing loss of reagent through the waste line. This is because the channels on the chip have a considerably smaller diameter (100 microns) compared to the inner diameter of the tubing (500 microns), and thus when this region is filled with liquid instead of air, the resistance of the overall fluid path is increased considerably. In fact, this increase in resistance is tunable based on the length and diameter of these channels, and on the chips we use in production settings, we modified the channel length to the waste outlet to increase resistance (and more importantly, increasing the resistance differential when filled with air vs. water) in order to further minimize loss of reagent in the Advance stage. For instance, on our platform, a 35uL fluid packet will take around 14s to travel from the rotary valve to the chip, whereas this same amount of fluid will take over 180s to flow through the waste line, and 600s to flow through the functional region of the chip. If the Advance stage is set at around 20s, on our workcell this guarantees that all of the air between the fluid packet and the valve controlling entry to the functional region of the chip will be cleared by the time the Introduce stage is reached (i.e. the packet reaches the chip), and this is true for all load volumes and reagents that we use on our platform. On the other hand, the additional 6s of flow through the chip's waste line produces a negligible waste volume, relative to the utilized volume from 10 minutes of active reagent introduction into the chip. Importantly, this ratio of waste to utilized volume is easily tunable. As you tune waste volume down, the duration of the Reset stage should increase. However, higher pressures can be used to decrease the Reset duration if this is of concern.

#### 1.3 LAIR robustness, independence of protocol steps, low dead volumes, and alternatives tested

The Advance and Reset stages can be set at durations that compensate for any variation in resistance that stems from differences in chip manufacturing, or from differences in upstream fluid paths, which may lead to small differences in loaded and cleared volumes. By resetting the fluid path to being filled entirely with air, the fluid handling for adjacent protocol steps is made independent and thus protocols become modular in a simple way, allowing one to for instance compose experiments by importing functions and calling those functions in the protocol file (e.g. we emphasized composability of experiments in Main Figure 5E). The liquid resting in the tubing between reagent source tubes and the rotary valve can also be conveniently returned to its source tube, which we do for instance when we choose to freeze an experiment in safe-mode in the event of an error like chip-clogging (Main Figure 4), or simply to cool this volume of reagent between successive flows of the same reagent. This allows us to drop from low dead volumes to near 0 dead volume when dealing with expensive samples. How this is achieved is similar to how cleaning cycles are executed after protocol runs depicted in Figure 3C.

We transitioned to the LAIR flow protocol from the field standard of using an inlet for each different reagent, which traditionally necessitates time-consuming and error-prone chip setup involving manual reagent-loading of many tubings (flow lines), more control lines, without control over reagent cooling. Our approach bypasses manual fluid handling entirely, since aliquots are simply screwed into the setup for use, and furthermore any dilutions are then generated automatically on-chip as opposed to manual off-chip serial dilutions. Even aliquots themselves are prepared for the freezer using a Hamilton Star liquid handling robot, to further standardize fluid handling and remove it from the hands of operators. Eliminating user participation through pipetting and reagent loading removes many possible sources of variability in an experiment. Our approach also allows us to achieve smaller flow volumes. For instance, whereas loading under 10$\mu$L of a reagent manually with a syringe can be challenging, we routinely use fluid packets of under 5uL with our workcell (resulting in under 5 nanolitres of reagent consumed per chamber).

LAIR is possible despite conventional wisdom in microfluidics for three reasons: (1) It does not matter if air travels on-chip in a region where a protein surface will not be analyzed (air must just be prevented from entering the functional region of the chip). (2) It does not take long to push a fluid packet to the chip relative to how long it takes to introduce this fluid into the chip. (3) Using a common-line where multiple different reagents pass is in general not a problem (as long as it is washed with PBS in between to clear reagent, and not re-used between experiments). This typically happens on-chip anyway (although to a lesser degree).

We evaluated alternatives to LAIR flow cycles, including strategies such as separating subsequent reagent packets from one another with bubbles or with PBS, but we determined that this results in accumulating volume error, and the delay between when a reagent is loaded and when it is being used on-chip complicates software control, makes subsequent reagent flows dependent on one another, and limits user-intervention in an experiment (which can be useful in the event of errors or in a research context).

##### 1.4 Pressure-based flowrate controller integrated with clogging-detection

Our flow-controller shows better performance than industry-standard syringe pumps, which may take up to 1 minute to reach flowrate setpoints, harbor 10% instability due to pulsing, and tend to be more constrained in their dynamic range[2]. This high degree of control is especially useful in sensitive applications, for instance in droplet microfluidics to improve monodispersity. Furthermore, the quick response is useful for exploring different flow regimes, for instance when tuning droplet diameter (as in Main Figure 5D). Importantly, compared to using a syringe pump, our system allows us to do real-time clogging detection (Main Figure 4), to cap pressures in order to prevent chip delamination, and is natively more compatible with Quake-valve based systems. Quake-valve systems like MITOMI[1,3,4,5] can possess different resistances depending on the number of unit-cell valves that are pressurized (e.g. the MITOMI button valve), or due to clogging, so a flowrate controller can improve the independence of protocol steps. However, if an inlet valve is shut or clogging occurs a syringe-pump will drive the chip to delamination.

#### 1.5 Leveraging chip-decoupling and on-chip multiplexing

With our standard in-house multiplexed chips (used for instance in Main Figure 5A), which consist of 32 rows and 32 columns, through multiplexing on-chip and chip decoupling we can run 96 (3 $X$ 32) experiment versions simultaneously. In this case, each experiment is executed across 32 different samples. Running multiple versions of experiments in parallel can be useful for making rapid experimental progress (for instance for assay development). Executing many experiments in parallel is made possible since the workcell already streamlines experiment design and execution. Furthermore, since we feed the reagents through two inlets to a given chip, the complexity of experiment setup does not scale as heavily with the number of reagents being used, as compared to standard setups that use 1 inlet per reagent. With the workcell, the additional reagent tubes need to just be screwed into hydra ports. For instance, typical assay development can involve months of iterated testing, learning, and redesigning. To develop assays more rapidly, it's possible to simply outline a large number of different assay versions that may or may not work (i.e. think through the conditionals of the typical outcome tree), leverage parallelization for execution, and if none of the designs produce a functional assay, then move on to the next research project.

#### 1.6 Automated cleaning circuit

We furthermore modified our platform to enable us to automate and standardize the cleaning of all reusable components, including the rotary valves, flowrate sensors, and the tubings between the pressure head and the rotary valves (reused if the same reagent is used subsequently). Through the way we configure the workcell, all of the internal fluid connections are made addressable by cleaning solution. At the beginning of an experiment, we screw in a cleaning solution (ethanol) into the pressure head, together with the other reagent tubes required for the experiment. A cleaning function in the software protocol file specifies a cleaning routine that is called at the end of every experiment. Cleaning solution is loaded into the output line between the first rotary valve and the chip. Then, the air pressure to the pressure head is shut off with the help of a solenoid valve (to create a pressure difference), and an air pressure line is created through the second rotary valve, into the chip inlet array, and backwards through the first rotary valve. The first rotary valve can then rotate through its different positions for protocol-specified durations in order to flush the previously loaded cleaning solution back through the rotary valve's internal channels and into its input tubings. Once the first rotary valve has cycled through all of its active positions, the role of the first and second rotary valves can be reversed, to clean the active ports in the second rotary valve. We only re-use input tubings where the exact same reagent will be used in the same position in the subsequent chip. We then also flush the flowrate sensor with cleaning solution to complete the cleaning routine.

#### 1.7 Image processing pipeline

For an accurate equilibrium binding assay, a measurement of the free antigen, bound antigen, and antibody levels at equilibrium was crucial. With these measurements, the occupancy level (the proportion of pulled-down antibodies occupied by an antigen, for 1:1 binding) could be established at different concentrations of free antigen, giving rise to a binding curve (Supplementary Figure 4B). In each chip chamber, a signal corresponding to the amount of molecules bound to the surface was measured in a circular region of interest (ROI) corresponding to the location of the button. The concentration of free antigen, on the other hand, was measured in a donut-shaped ROI corresponding to the chamber excluding the button. Since the chip fabrication process sometimes resulted in a small misalignment between the buttons and chambers, it was crucial to detect both ROIs independently of each other. This ROI detection was done using a brightfield scan (Supplementary Figure 4A).

To detect the chamber, a high-pass filter of kernel size 3 pixels was first applied to each microscope image (Supplementary 4C, top). In the resulting image, diagonal lines were detected using two diagonal filters with a kernel size of 12 $\mu$m (one filter for each diagonal direction) and a subsequent threshold. The largest object in each quadrant then corresponded to the edge of the chamber, so the chamber could be identified as the largest inner circle within these edges. If the radius of the chamber did not fall into a predefined range, the process was re-iterated with an increased kernel size of the diagonal filters. After the chamber, detection of the button ROI started with applying a Li threshold on all pixels inside the chamber (Supplementary Figure 4C, bottom). The resulting image was divided into radial bins, and in each bin the point closest to the chamber centre was kept. The button was identified as the largest inner circle contained within these points. If the detected button radius did not fall within a predefined range, the process was re-iterated using an increased threshold in the first step. Based on this button detection, the button ROI was defined as a circle with a radius of 27 $\mu$m with its centre corresponding to the centre of the button detection. The chamber ROI corresponded to the chamber detection with an inner circle (with radius 41 $\mu$m and its centre at the centre of the button) removed.

To quantify fluorescence intensities, the background of each fluorescence scan was removed using a separate background scan taken at the start of the experiment. The mean of the 80th-interpercentile range of all background-subtracted pixels in a ROI was calculated. Occupancy levels were calculated by dividing the mean fluorescence of the bound antigen (or solution binding partner) by that of the bound antibody (or binding partner on the surface) in each button ROI. The level of free antigen was the mean of the chamber ROI, which was converted to a concentration using a calibration curve.

The algorithm was implemented as software on Amazon Web Services (AWS). Microscopy images were uploaded to Amazon's cloud object storage (S3) service, together with a YAML file specifying (1) the set-up of the experiment and image filenames and (2) the ROI-detection parameters. The upload of the YAML file triggered an AWS Lambda function which submitted each computation as an AWS Batch Job (Figure 5). When the ROI-detection job was finished, the ROI quantification jobs were initiated automatically (one job for each fluorescence scan). Finally, the images of each scan were stitched into a whole-chip overview for manual inspection, and the results were automatically uploaded to Notion.

### References


[1] S. J. Maerkl and S. R. Quake, "A systems approach to measuring the binding energy landscapes of transcription factors," *Science*, vol. 315, pp. 233–237, Jan. 2007.

[2] Fluigent, "Microfluidic droplet generation using syringe pumps and pressure-based flow controllers." `https://www.fluigent.com/wp-content/uploads/2022/01/microfluidic-droplet-generation-using-different-flow-controllers.pdf`, 2019. Accessed: 2023-01-10.

[3] P. M. Fordyce, D. Gerber, D. Tran, J. Zheng, H. Li, J. L. DeRisi, and S. R. Quake, "De novo identification and biophysical characterization of transcription-factor binding sites with microfluidic affinity analysis," *Nature biotechnology*, vol. 28, no. 9, pp. 970–975, 2010.

[4] A. K. Aditham, C. J. Markin, D. A. Mokhtari, N. DelRosso, and P. M. Fordyce, "High-

throughput affinity measurements of transcription factor and DNA mutations reveal affinity and specificity determinants," *Cell Systems*, vol. 12, no. 2, pp. 112–127, 2021.

[5] C. Markin, D. Mokhtari, F. Sunden, M. Appel, E. Akiva, S. Longwell, C. Sabatti, D. Herschlag, and P. Fordyce, "Revealing enzyme functional architecture via high-throughput microfluidic enzyme kinetics," *Science*, vol. 373, no. 6553, p. eabf8761, 2021.
